# Computational and Experimental Evaluation of a Flow-Conditioning Anastomotic Device for Arteriovenous Fistula Maturation

**DOI:** 10.64898/2026.02.06.704514

**Authors:** Keith Saum, Begoña Campos, Diego Celdran Bonafonte, Liran Oren, A. Phillip Owens, Prabir Roy-Chaudhury

**Author notes:** corresponding author: Keith Saum, University of Michigan, 5313 Brehm Center, 1000 Wall St, Ann Arbor, MI 48105, USA. Phone: (734) 936-***.

## Abstract

The arteriovenous fistula (AVF) is the preferred method of vascular access for hemodialysis; however, 30-50% of AVFs undergo primary failure and are unsuitable for clinical use. As disturbed hemodynamics initiate endothelial injury and intimal hyperplasia, we designed an endovascular flow-conditioning anastomotic device (FCAD) to directly improve AVF hemodynamics and protect the anastomotic region. Using computational fluid dynamics, we characterized the flow field and wall shear stress (WSS) profiles in idealized AVF models with and without the FCAD. Incorporation of the FCAD into a brachiocephalic AVF model reduced regions of oscillatory WSS and generated a symmetrical flow profile in the draining vein compared to a reference AVF. Parametric studies also identified an FCAD geometry with a tab angle, height, and aspect ratio of 30°, 0.1 diameters, and 1.0 restored time-averaged WSS along the inner venous wall, achieving a physiological level without inducing regions of oscillatory flow throughout the cardiac cycle. Similar findings were observed with an in vitro model using particle imaging velocimetry. This study demonstrates the feasibility of the FCAD to normalize venous flow and WSS while imposing minimal resistance to blood flow. Restoring physiological WSS levels on the venous wall is expected to preserve endothelial function and improve AVF maturation.

## Introduction

The arteriovenous fistula (AVF) is the preferred form of vascular access for hemodialysis; however, 25% to 60% of AVFs fail to mature or develop an adequate blood flow or luminal diameter to support dialysis.^1^ AVF maturation failure results from intimal hyperplasia (IH) and lack of outward remodeling, leading to progressive occlusion of the venous anastomosis and the draining vein.^2^ While the exact mechanisms underlying AVF maturation failure are unclear, considerable evidence suggests that the disturbed hemodynamics within an AVF play an important role in IH formation. Specifically, regions of oscillatory flow with low wall shear stress (WSS) appear to correlate with sites vulnerable to IH formation.^3^ These disturbed hemodynamics may alter local endothelial cell function and initiate the development of IH.^4^ In light of this understanding, hemodialysis could benefit from the development of a novel vascular access technology capable of restoring physiological hemodynamics within the AVF and draining vein.

The degree of oscillatory WSS downstream of the AVF anastomosis could be potentially reduced using flow-conditioning technology. Flow conditioning is a method used to alter the flow profile of a fluid downstream of a disturbance or obstruction.^5^ Flow-conditioning devices consist of mechanical damping elements inserted into the lumen of a conduit, such as honeycomb vanes, perforated plates, tabs, and tube bundles. In particular, tab-type flow conditioners have proven most effective for particulate mixtures due to the tab’s tapered design, which minimizes fouling and pressure losses.^5,6^ In addition, they are the only geometry that can be retrofitted into elbows, similar to an AVF configuration. For these reasons, we hypothesized that incorporating tab-type flow-conditioning elements into an endovascular device implanted at the AVF anastomosis would reduce regions of low, oscillatory WSS along the outflow vein of a fistula without imposing significant resistance to AVF blood flow.

In an attempt to minimize WSS abnormalities and adverse venous remodeling, several studies have attempted to identify an optimal anatomical configuration for an AVF.^7,8^ However, as concluded by others, it is very difficult to identify an AVF configuration that abolishes all areas of disturbed WSS.^3^ Rather, an important goal for improving AVF maturation could be the combination of a “reasonable” configuration, together with a means of reducing disturbed WSS in the vein. We have developed a novel flow-conditioning anastomotic device (FCAD) to improve AVF maturation by normalizing venous flow and protecting the anastomotic region.^9^ The endovascular device is implanted into the AVF at the time of creation and consists of arterial and venous lumens coupled to mimic an end-to-side AVF anastomosis (Fig. 1A). The venous lumen of the FCAD features flow-conditioning tabs along the outer curve and outlet of the swing segment, which are responsible for restoring a physiological flow and WSS profile at the venous outlet (Fig. 1B-C). The clinical implementation of this technology would consist of implanting the arterial and venous segment into the respective lumens prior to anastomosis creation (Fig. 1D). This endovascular approach also allows for standardization of the AVF configuration and interventional techniques such as thrombolysis and angioplasty to still be performed in the advent of clot formation. As a first step toward the development of this new technology, the objectives of this study were to computationally characterize the flow and WSS profiles of a FCAD compared to an idealized AVF and to investigate the effects of the tab geometry on flow characteristics.

**Figure 1:**
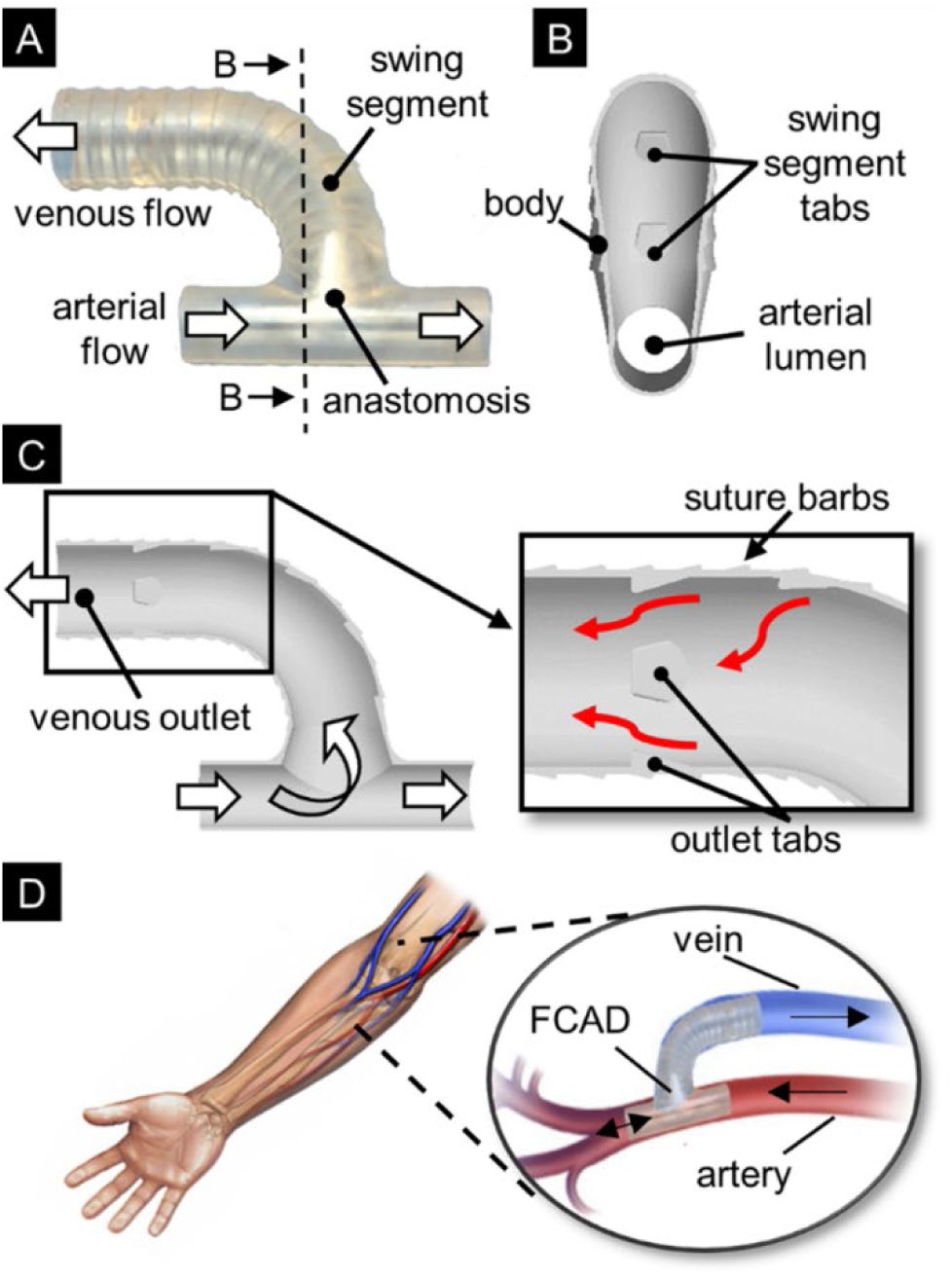
FCAD design and operation principle. [**A**] Prototype FCAD featuring a 90° anastomotic venous swing segment. Arrows indicate the direction of blood flow through the device. [**B**] Transverse section through the anastomosis showing flow-conditioning tabs along the venous swing segment. [**C**] Longitudinal section through the FCAD lumen showing a second set of flow-conditioning tabs at the venous outlet. Red arrows indicate direction of blood flow around the tabs. [**D**] Clinical implementation during AVF creation. Black arrows indicate the main direction of blood flow.

## Materials and Methods

### Idealized AVF Models with the FCAD

Idealized models of an end-to-side brachiocephalic AVF with and without the FCAD implant were generated using SolidWorks 2015 (Dassault systems, Velizy-Villacoublay, France). We generalized vessel shape, assuming that each patient exhibits a unique vessel size and shape. The artery and vein was modeled as rigid circular cylinders with vessel diameters of 4.0 and 6.0 mm, respectively, with an anastomotic angle of 90°. The length of the proximal artery (PA), distal artery (DA), and vein were 12 times the vessel diameter in order to have sufficient hydraulic length to establish fully-developed flow and to characterize the flow field in the vein downstream of the anastomosis (Fig. 2A). The swing segment of the venous branch was generated with a radius of curvature equal to twice the venous diameter, and the juxta-anastomotic region of the vein was tapered for a length of two diameters to ensure a smooth transition between the artery and venous sections. For the FCAD implant models, we generated flow-conditioning tabs along the swing segment of the vein and outlet of the venous elbow (Fig. 2B). To characterize the impact of the tab geometry on the venous flow field, FCAD implant models were generated with four tab angles (15°, 30°, 45°, and 60°), four tab heights normalized to the venous diameter (0.1D, 0.2D, 0.3D, and 0.4D), and four tab aspect ratios (0.75, 1.0, 1.25, and 1.5) as shown in Figure 2C. A matrix of the FCAD models considered in the parametric analysis is shown in Table 1.

**Figure 2:**
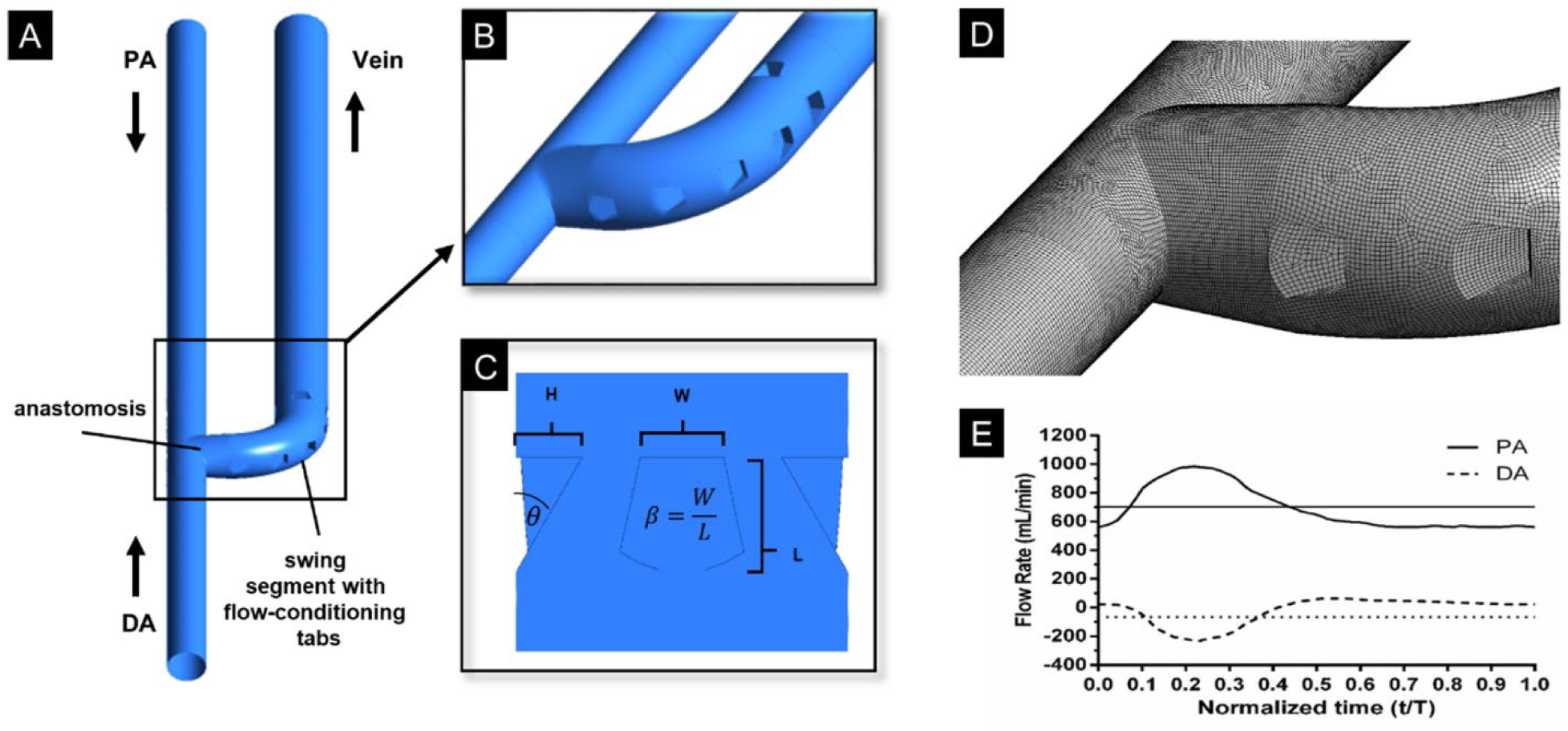
Idealized AVF model with FCAD geometry for CFD simulations. [**A**] 3D surface of the idealized AVF. Black arrows indicate the direction of blood flow in the proximal artery (PA), distal artery (DA), and venous outlet. [**B**] Detailed view of the flow conditioning tabs along the venous swing segment and outlet. [**C**] Detailed view of the tab dimensions at the venous outlet used in the parametric analysis including the tab angle (θ), height (H), and aspect ratio (β). [**D**] Surface mesh at the anastomotic region. [**E**] Volumetric flow rate waveforms of a brachiocephalic AVF obtained from He et al. Continuous and dashed curves represent blood flow rate in PA and DA, respectively. Blood flow in the DA changes direction during the cardiac cycle; negative flow is antegrade (towards the hand) and positive flow is retrograde (towards the anastomosis). Horizontal lines indicate time-averaged blood flow rate over the entire cardiac cycle, 705 mL/min for PA and 66 mL/min for DA, respectively.

**Table 1:**
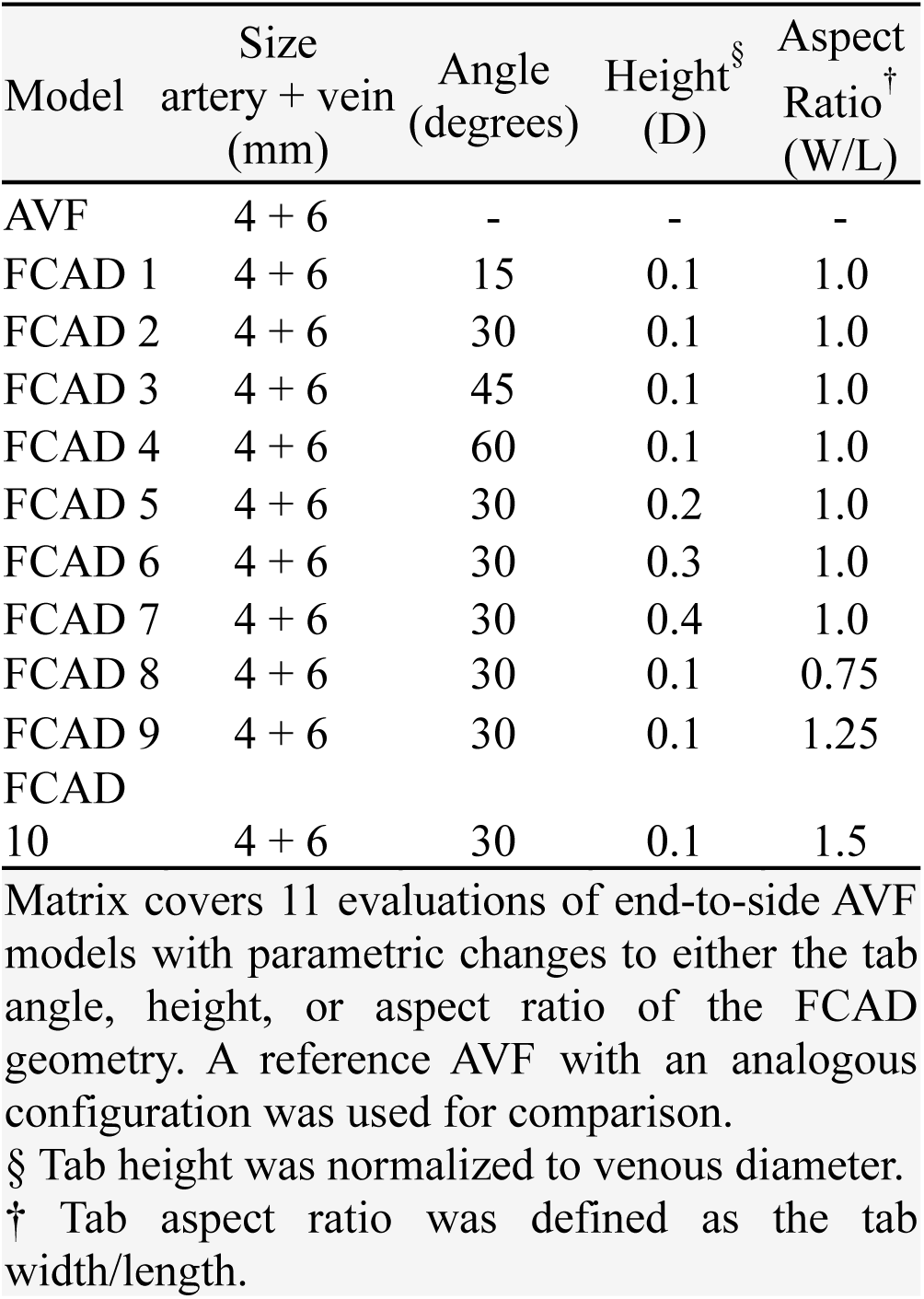
Matrix of Parametric FCAD Models

The fluid volume of each model was discretized with a structured hexahedral mesh generated in ICEM 16.0 (ANSYS, Inc., Canonsburg, PA). A boundary layer of 5 cells with a growth rate of 1.2 was generated at the walls to accurately capture changes in WSS. Three meshes of increasing element density were generated for a single model, with and without the FCAD geometry, to estimate discretization error using the grid convergence index (GCI) method.^10^ Steady-state computational fluid dynamics (CFD) simulations, corresponding to the venous flow rate at peak systole, were performed in each set of three meshes and used to estimate the mean velocity and mean WSS in the flow domain. The results of the grid convergence study are presented in Supplemental Table 1. Velocity and WSS fluctuations due to poor spatial resolution were minimized to less than 5% when the mesh size exceeded 5.4 × 10^6^ and 7.0×10^6^ elements in the AVF and FCAD models, respectively. Therefore, we generated similar meshes (∼7.0 × 10^6) for all remaining FCAD models.

### CFD Simulations of Blood Flow

Idealized AVF models were imported into the commercial CFD solver ANSYS Fluent 16.0 (Fluent, Inc., Lebanon, NH) and used to simulate pulsatile blood flow in the vessels. The transient Navier–Stokes equations were solved using a laminar model with a pressure-implicit with splitting of operators (PISO) algorithm for the pressure–velocity coupling. As boundary conditions, volumetric flow waveforms were prescribed at the PA and DA inlets (Fig. 2E), obtained from He et al. by 2D cine PC MRI of an end-to-side brachiocephalic AVF 5 months after creation.^11^ A traction-free boundary condition was applied at the venous outlet, and zero velocity (no-slip condition) was applied at the vessel walls, which were considered to be rigid. Time-dependent terms were discretized in an implicit scheme with second-order accuracy. A total of 1,000 fixed time steps were used per pulse cycle, yielding a time step size of 0.001 s and a cardiac cycle period of 1 s. For each simulation, three complete cardiac cycles were solved to avoid start-up transients, and only the third cycle was saved for data processing in 100 equal time steps. A residual of 10^−5^ was set as the convergence criterion. Blood was modeled as an incompressible fluid with a constant density of 1.050 g/cm^3^, and blood viscosity was considered non-Newtonian by using the Carreau rheological model, described previously as:

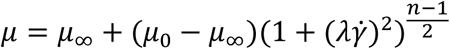

where 𝜇_∞_ is the limiting viscosity at infinite shear rate, 𝜇_0_ is the limiting viscosity at zero shear rate, λ and n are constants, and 𝛾̇ is the strain rate.^12^ The use of the Carreau model was justified by our focus on identifying venous wall regions with low WSS, for which non-Newtonian effects may be particularly significant. For the PA inlet and venous outlet, we also calculated the Reynolds and the Womersley numbers as described previously.^13^ Geometric and hemodynamic features of the AVF and FCAD model 2 simulations are summarized in Supplemental Table 2. The mean and range of the Reynolds numbers predicted at the vein outlet were 1315 (1137–2228) and 1643 (1343-2382) for the AVF and FCAD, respectively, which justifies the use of a laminar flow solver.

To investigate the resistance to AVF flow imposed by FCAD implantation, we also performed CFD simulations on each AVF model by scaling the PA and DA waveforms to generate a series of flow regimes with increasing Reynolds number. These studies were used to calculate the pressure-loss coefficient of the FCAD across a range of flow regimes in which the device would operate in vivo. Further details of these studies are available in the online Electronic Supplemental Materials.

### Flow Field Characterization and Metric Analysis

The ability of the FCAD to normalize venous AVF flow and WSS was investigated *in silico* by comparing flow profiles and WSS characteristics in the idealized AVF model (i.e., reference hemodynamic state) with those in FCAD AVF models subjected to parametric changes in their tab geometry. Once the flow field of each AVF was obtained, 13 transverse cross-sections were created normal to the centerline of the vein at 6 mm increments, starting at the anastomosis. The flow profile in each cross-section was assessed using velocity contours and the following absolute and relative efficiency parameters, as previously described in the literature.^6^ By comparing these parameters at multiple locations downstream of the FCAD outlet to the corresponding location in the reference AVF model, we were able to tailor the performance of the device to improve venous flow through the AVF.

The profile symmetry, K_sym_, was used to assess the symmetry of the time-averaged velocity profile about the vessel centerline. This parameter represents a non-dimensional distance between the centroid of the flow profile and the vessel axis and can be calculated in any transverse cross-section of the vessel and is expressed as:

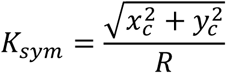

where R is the vessel radius, and x_c_ and y_c_ are the coordinates of the centroid of the mass flow given by:

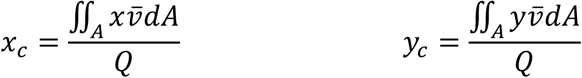

where v is the time-averaged axial velocity, Q is the volumetric flow rate, and x and y are distances from the centerline to the radial coordinate. Therefore, K_sym_ is always positive, with smaller values indicating a less distorted flow.

Changes in WSS through the FCAD and downstream of the venous outlet were characterized using two WSS metrics to assess regions of potential endothelial dysfunction and/or IH formation. Time-averaged WSS (TAWSS) was used to identify regions of excessively high or low WSS that occur over the cardiac cycle and was defined as:

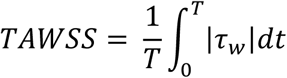

where T is the period of the cardiac cycle, and τ_w_ is the instantaneous WSS vector. To quantify reciprocating disturbed flow during the cardiac cycle, the oscillatory shear index (OSI) was calculated on the surface of each AVF model as previously described^14^:

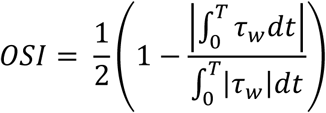

These WSS parameters were also averaged circumferentially within each of the 13 cross-sections, as previously described, yielding a single value for each parameter at multiple intervals along the AVF vessel.^11^

On the basis of the absolute parameters presented above, we also determined the relative efficiency of the FCAD at improving each flow-field metric, ε_i_, compared to the reference AVF model as:

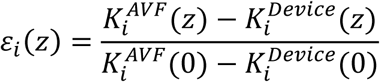

where K^AVF^ and K^Device^ represent the values of the absolute parameters calculated for the same vessel configuration without and with FCAD, respectively, and z is the distance downstream of the AVF anastomosis. Therefore, these parameters quantify the efficiency of the FCAD relative to the reference AVF as the distance downstream of the implant varies. Values of ε_i_ greater than zero show an efficiency of the FCAD greater than the reference AVF and allow for comparison of different FCAD geometries. Further details on the flow-field characterization are provided in the online Electronic Supplemental Materials.

### Fabrication of AVF flow phantoms

A two-step, embedded scaffold removing open technology (ESCARGOT)^15^ method was used to generate AVF flow phantoms for assessing the FCAD performance in vitro using time-resolved particle imaging velocimetry (PIV). Briefly, 3D models of the vessel lumens from idealized AVF models with and without the implant design, described above, were extracted using SolidWorks 2015 (Dassault Systems, Velizy-Villacoublay, France). These CAD models were converted into STL format to generate artificial vessels by 3D printing. The artificial vessels were 3D printed in acrylonitrile butadiene styrene (ABS) plastic using a commercial 3D printer (Ultimaker 2+, Ultimaker; Geldermalsen, Netherlands) with a 0.15mm nozzle (3D Solex, model #7072482000109) at the highest layer resolution mode (60 μm). During post-processing, a small amount of acetone was used to smooth any rough surfaces. Each 3D ABS model was then encapsulated in transparent polydimethylsiloxane (PDMS) (Sylgard 184, Dow Corning; Midland, MI) and cured under vacuum at room temperature overnight. After curing, the PDMS blocks containing the 3D ABS model were cut into a rectangular shape. Subsequently, the blocks were placed in an acetone bath overnight to dissolve the ABS, resulting in a hollow AVF flow phantom for each model. Each phantom was then rinsed with water and allowed to dry before coating the luminal surface with PEG-200 to reduce particle sticking on the channel during PIV experiments. Supplemental Figure 1 shows example images of this fabrication process after 3D printing, embedding, and removal of the artificial AVF vessels.

### PIV setup and analysis

The AVF phantoms were incorporated into an in vitro flow system and perfused with a working fluid composed of water (55%) and glycerol (45%) to match the refractive index of transparent PDMS. Supplemental Figure 1D-F demonstrates the refractive index matching of this working fluid compared to water and air. Due to the increased viscosity of the working fluid, the flow waveforms shown in Figure 2E were scaled to match the Reynolds numbers at the PA inlet and PA outlets, respectively. A pulsatile blood pump (55-3321, Harvard Apparatus, MA) was used to propel the working fluid into the PA of the phantom at a rate of 60 beats per minute, corresponding to a mean Womersley number of 3.4. An adjustable-height afterload was connected to the DA outlet of the phantom to achieve the desired flow ratio between the DA and vein throughout the pump cycle. The venous outlet was left free. Flow rates at the PA inlet and DA outlet were monitored throughout the experiments using two perivascular flow probes (4PS and 6PS, Transonic Systems Inc., Ithaca, NY) connected to a Transonic T402 flowmeter console. The pressure at the PA inlet was also monitored using an inline pressure transducer (PRESS-N-025, PendoTECH; Princeton, NJ). Acquisition of the flow and pressure transducer signals was made at 1 kHz and synchronized with the PIV measurements using a data acquisition system (Powerlab, AD Instruments; Colorado Springs, CO). A schematic of the flow system setup is shown in Supplemental Figure 2.

Flow-velocity measurements in each phantom were obtained using time-resolved 2-D PIV. The working fluid was seeded with 2.0 µm polystyrene particles coated with Rhodamine to capture the flow field. PIV images were acquired in the mid- to longitudinal plane of the vessels by illuminating the flow with a high-repetition-rate, dual-cavity Nd:YLF laser system (LDY304, Litron; Agawam, MA) synchronized with a high-speed video camera (Phantom 1610, Vision Research Inc.; Wayne, NJ). The laser was used to generate a 1 mm-thick light sheet in the anastomosis region near the outflow venous segment. The mid-longitudinal plane was chosen because vessel area and velocity magnitudes are typically highest in this section, and it enabled visualization of the FCAD geometry. The camera lens was fitted with a 527nm band-pass filter to minimize reflections of the laser. An area of 60 × 40 mm in physical space was captured for each image, corresponding to a pixel resolution of 1280 × 800. The PIV data was captured at 200 Hz, and each image pair was taken at a time interval of 150 μsec. Post-processing of the PIV data was performed using DAVIS 8.4 (LaVision GmbH, Goettingen, Germany) with a multi-pass, decreasing window size (64 × 64 to 32 × 32) and an adaptive interrogation window with 50% overlap. A total of 6000 image sets were acquired over 30 pump cycles and phase-averaged. The phase of the velocity fields in relation to the pump cycle was determined by synchronizing the PIV measurements with the pressure transducer signal.

## Results

### Flow Patterns in the AVF and FCAD

CFD-derived velocity contours of blood flow in the longitudinal symmetry plane of the idealized AVF and an FCAD-implanted AVF are shown in Figure 3 at peak systole, diastole, and time-averaged across the entire cardiac cycle. Representative velocity profiles orthogonal to the vessel centerlines are also shown at the venous outlet, mid-anastomosis, and arterial outlet of the FCAD in both models. In the AVF, the high-momentum flow emanating from the PA is deflected towards the outer curve of the venous swing segment, resulting in a distorted and asymmetric flow profile in the draining vein throughout the cardiac cycle (Fig. 3A – 3C). This flow pattern creates a region of flow stagnation and recirculation along the inner curve of the venous swing segment and draining vein, which is accentuated during the diastolic phase of the cardiac cycle. In addition, the sudden curvature of the venous swing segment of the AVF leads to the formation of Dean vortices characteristic of curved tubes. These vortices are observed in the orthogonal cross-section at the venous outlet in Figure 3A and gradually diminish along the length of the vein.

**Figure 3:**
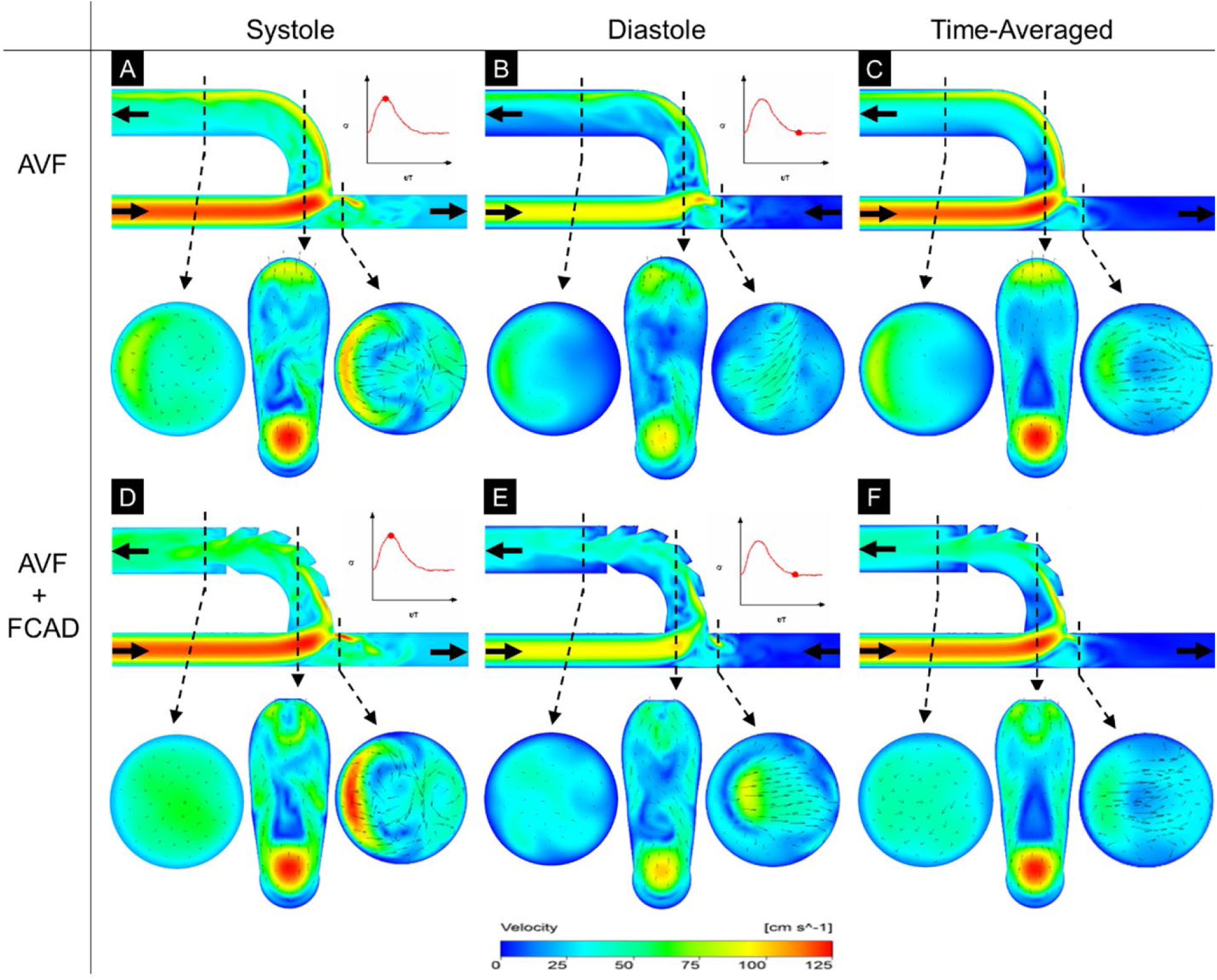
Flow field profiles in the AVF and FCAD model 2. Temporal snapshots of velocity magnitude on the longitudinal symmetry plane in idealized end-to-side AVF configurations with and without FCAD 2 implanted. Velocity contours are shown at peak systole (**A** & **D**), mid-diastole (**B** & **E**), and time-averaged across the entire cardiac cycle (**C** & **F**). Velocity vectors on three planes orthogonal to the vessel centerlines are shown below each time-point, positioned at the venous outlet, mid-anastomosis, and arterial outlet of the FCAD. Solid black arrows indicate the direction of blood flow.

In contrast, implantation of an FCAD with a tab angle of 30° and a height of 0.1 diameters re-directs blood flow from the outer wall of the swing segment back toward the centerline of the vein (Fig. 3D-F). After passing through the flow-conditioning tabs along the venous outlet of the FCAD, the region of flow stagnation along the inner curve of the draining vein is significantly reduced, and the velocity profile appears more symmetric in the longitudinal and orthogonal planes compared to the AVF. These flow characteristics persist throughout the entire cardiac cycle. Flow through the DA outlet of the FCAD is almost identical to the AVF due to impingement of flow on the toe of the anastomosis, resulting in a region of flow stagnation on the anastomosis floor.

### WSS Patterns in the AVF and FCAD

Given the relationship between regions of low, oscillatory WSS and IH formation in the venous limb of native fistulae, we assessed changes in WSS parameters in AVF models with and without the FCAD. Time-averaged flow streamlines and the corresponding TAWSS over the vessel surface in the AVF and an FCAD model 2 are shown in Figure 4. The regional TAWSS distribution on the venous wall of the AVF (Fig. 4B) revealed a region of locally low WSS along the inner wall of the AVF vein proximal to the anastomosis. In contrast, this region was absent in the FCAD-implanted AVF with a tab angle of 30° and a height of 0.1 diameters. However, small regions of low WSS were identified downstream of each flow-conditioning tab within the FCAD lumen. Analysis of the OSI between each model demonstrated similar patterns of highly oscillatory flow (OSI > 0.4) at the anastomosis floor and the just-anastomotic vein (Fig. 4C). In addition, small regions OSI > 0.4 were also present around the perimeter of each flow-conditioning tab. Importantly, regions of oscillatory flow (OSI > 0.001) at the venous outlet were significantly smaller in the FCAD model 2 than in the reference AVF, particularly along the inner wall with low TAWSS.

**Figure 4:**
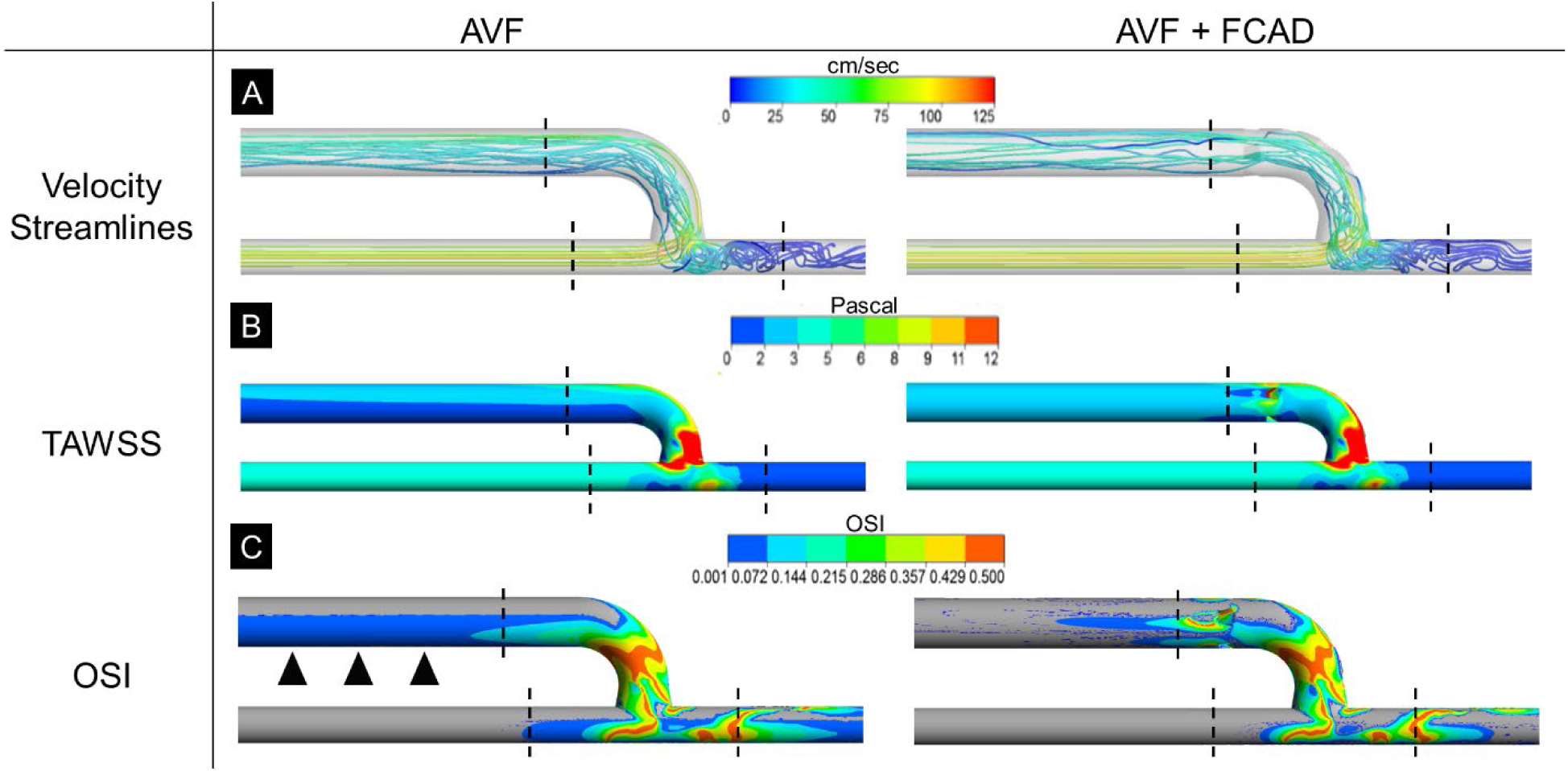
Wall shear stress analysis in AVF and FCAD model 2. [**A**] Time-averaged velocity streamlines in the reference AVF and FCAD model 2. [**B**] Time-averaged wall shear stress (TAWSS) in the reference AVF and FCAD model 2. [**C**] Oscillatory shear index (OSI) in the reference AVF and FCAD model 2. Black arrowheads highlight the difference in OSI along the inner wall of the venous outlet between the AVF and FCAD. Vertical dashed lines represent the corresponding location of the FCAD inlets and outlets in each view.

### Effect of FCAD Design on AVF WSS and Flow Field

To investigate the impact of the tab design on flow characteristics more closely, we circumferentially averaged the OSI at multiple locations downstream of the anastomosis to compare WSS between the idealized AVF model and multiple FCAD models with parametric variations in tab angle, height, and aspect ratio. In the idealized AVF, mean OSI was highly skewed along the venous swing segment with a mean value of 0.225 and decayed exponentially downstream (labeled “AVF” in Fig. 5A-C). Incorporation of an FCAD with tabs having an angle of 30-60°, a height of 0.1-0.2 diameters, or an aspect ratio of 0.75-1.00 significantly improved mean OSI downstream of the venous outlet (Fig. 5A-C). In contrast, extreme values of the tab height and aspect ratio worsened the mean OSI downstream of the venous outlet. Interestingly, the mean OSI in the region adjacent to FCAD implant (between the anastomosis and venous outlet) was significantly higher for most tab geometries but decayed more rapidly than the reference AVF. Calculation of the relative efficiency at improving the mean OSI, ε_OSI_, for each design compared to the reference AVF revealed that a tab angle, height, and aspect ratio of 30°, 0.1 diameters, and 1.0 resulted in the lowest mean OSI downstream of the FCAD, respectively (Fig. 5D – F).

**Figure 5:**
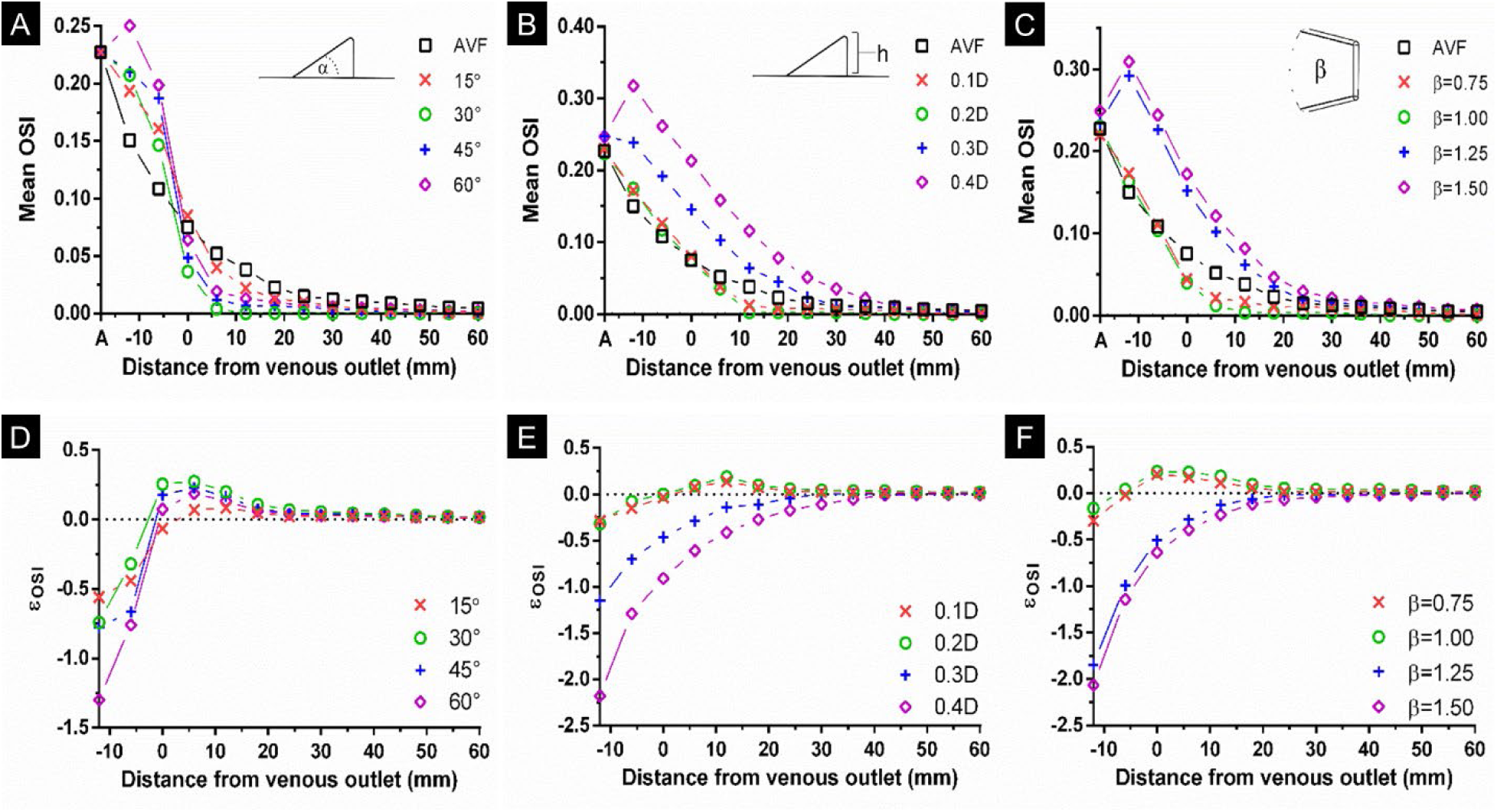
Parametric analysis of oscillatory WSS along the draining vein in AVF and FCAD models. Oscillatory shear index (OSI) was circumferentially averaged and plotted at multiple cross-sections downstream of the anastomosis in FCADs with increasing [**A**] tab angle (α), [**B**] tab height, and [**C**] tab surface area ratio. The location of the FCAD venous outlet was set to be 0, and the anastomosis is labeled “A” in each plot axis. The reference AVF without an FCAD is labeled “AVF”. The relative efficiency (ε) of each FCAD at improving mean OSI compared to the reference AVF was also determined for each tab angle (**D**), height (**E**), and area ratio (**F**).

In addition, we also used the time-averaged velocity fields to compare the flow profile symmetry, K_sym_, between the idealized AVF model and multiple FCADs at multiple locations downstream of the venous outlet (Supplemental Fig. 3). In the idealized AVF, velocity profile symmetry was highly skewed at the location corresponding to the venous outlet of the FCAD with a symmetry number value of 0.49, and decayed exponentially downstream (labeled “AVF” in Supplemental Fig. 3A-C). Incorporation of an FCAD with tabs having an angle of 30-45°, a height of 0.1-0.2 diameters, or an aspect ratio of 0.75-1.25 significantly improved profile symmetry at the venous outlet. In contrast, tabs with an angle of 15° had little effect on the flow profile. Calculation of the relative efficiency of the profile symmetry number, ε_Ksym_, relative to the reference AVF revealed that a tab angle, height, and aspect ratio of 30°, 0.1 diameters, and 1.0, respectively, also yielded the most symmetrical flow profile (Supplemental Fig. 3D-F). Conversely, a tab height > 0.2 diameters or an aspect ratio > 1.25 resulted in a more distorted flow profile compared to the idealized AVF, due to the high degree of obstruction.

### Generation of Counter-Rotating Vortices by FCAD Tabs

Figure 6 shows longitudinal and isometric views of local normalized helicity (LNH) isosurfaces at peak-systole and mid-diastole in AVF models with and without the FCAD model 2. For reference, velocity streamlines are shown next to each isometric view. Coherent helical flow structures originate at the anastomosis in the AVF with both clockwise and counterclockwise rotation, as demonstrated by the blue and red color isosurfaces in the upper and lower surfaces of the venous swing segment, respectively (Fig. 6A and C). These helical structures are present throughout the cardiac cycle, with the highest prevalence during systole. In FCAD AVF models, the generation of counter-rotating vertical structures is observed along the trailing edge of each tab, demonstrated by the reciprocal change in LNH isosurface color highlighted by black arrowheads (Fig. 6B and D).

**Figure 6:**
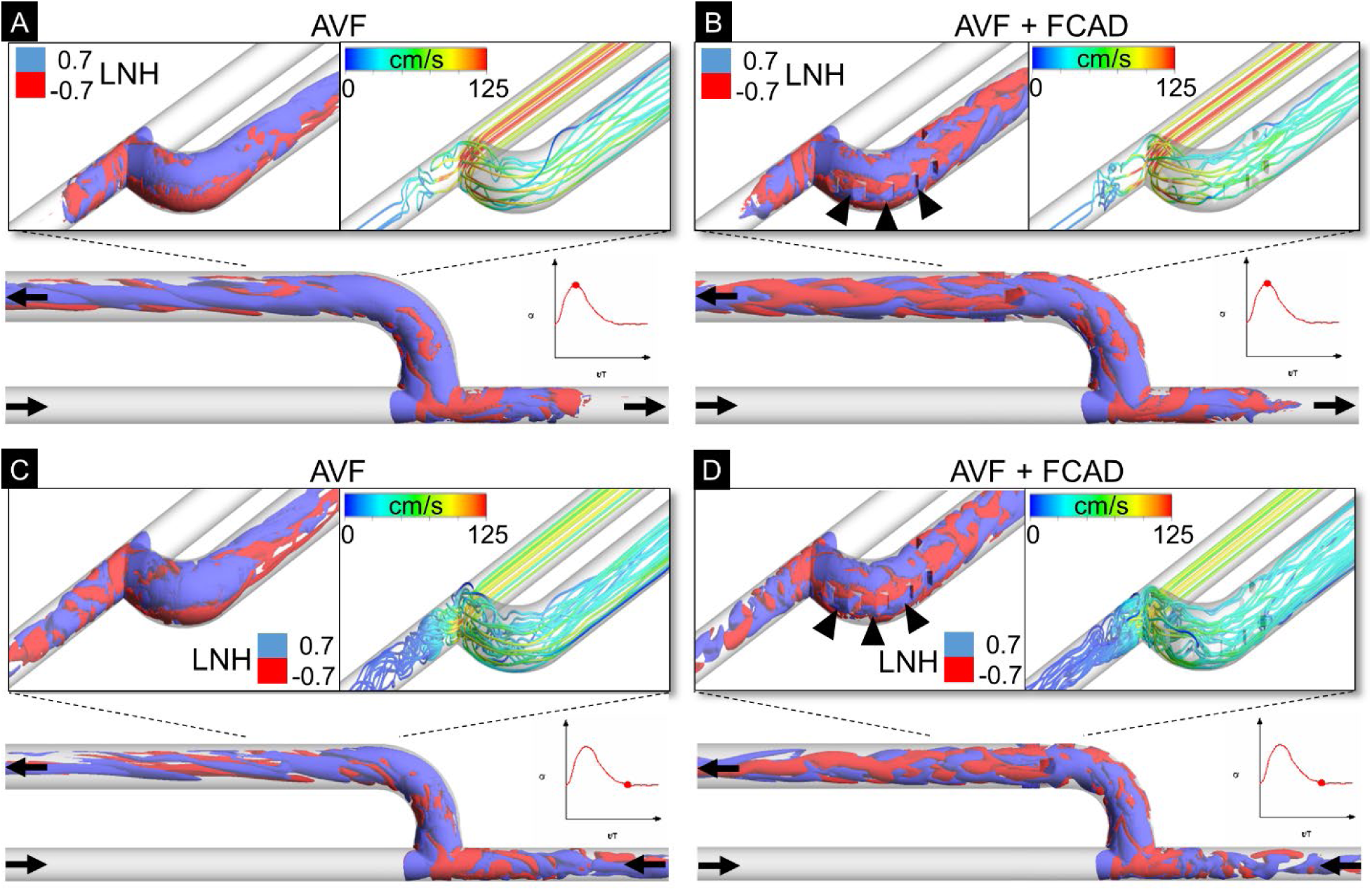
LNH isosurfaces in the AVF and FCAD model 2. Isometric and longitudinal views of local normalized helicity (LNH) isosurfaces along the venous swing segment in idealized end-to-side AVF configurations with and without FCAD 2. LNH isosurfaces are shown at peak systole [**A & B**] and mid-diastole [**C & D**]. Threshold values of LNH (±0.7) are used for the identification of clockwise and counter-clockwise rotating helical structures. Velocity streamlines are shown next to each isometric view for reference. Arrowheads show the generation of counter-rotating helical structures at the trailing edge of each flow-conditioning tab. Solid black arrows indicate the direction of blood flow.

### FCAD Imposes Minimal Resistance to AVF Blood Flow

Resistance to blood flow imposed by implantation of the FCAD was assessed by calculation of the pressure loss coefficient for each model across a series of flow regimes with increasing Reynolds number (Supplemental Fig. 4). In the AVF, the pressure loss coefficient was the greatest (2.65) at the lowest Reynolds number assessed (Re = 760) and decayed exponentially with increasing Re. Incorporation of an FCAD with tabs having an angle of 15-30°; a height of 0.1-0.2 diameters; or an aspect ratio of 0.75 – 1.0 had minimal impact on the pressure loss coefficient, as demonstrated by the overlapping line in Supplemental Figures 4A – C. In addition, the mean pressure loss across the anastomosis was similar between the AVF and FCAD designs at 66 (52 – 78) and 69 (53 – 81) mmHg for AVF and FCAD model 2, respectively (Supplemental Table 2). These results suggest that the FCAD geometry imposes minimal resistance to AVF flow rate. When combined with the flow-field data, tabs with an angle of 30°, a height of 0.1 diameters, and an aspect ratio of 1.0 produced the most symmetric venous flow profile, with minimal oscillatory WSS and minimal resistance to flow.

### PIV Assessment of AVF and FCAD Flow

To validate the performance of the FCAD in our CFD studies, we utilized PIV to quantify velocity flow fields in idealized AVF flow phantoms with and without the FCAD geometry. PIV-derived velocity fields in the longitudinal symmetry plane of the idealized AVF and FCAD phantoms are shown in Figure 7 at peak systole, diastole, and time-averaged across 30 flow cycles. Consistent with our CFD findings, the high-momentum flow emanating from the PA is deflected toward the outer curve of the venous swing segment in the AVF model, resulting in a distorted and asymmetric flow profile in the draining vein throughout the cardiac cycle (Fig. 7A & C). In addition, the region of flow stagnation and recirculation along the inner curve of the venous swing segment and draining vein was also observed. In contrast, the FCAD phantom with a tab angle of 30° and a height of 0.1 diameters redirects the fluid flow from the outer wall of the swing segment back toward the centerline of the vein, resulting in a fairly symmetric flow field in the downstream vein. These flow characteristics also persist throughout the entire cardiac cycle. Further similarities between the numerical CFD simulations and the PIV studies are shown in Supplemental Figure 5.

**Figure 7:**
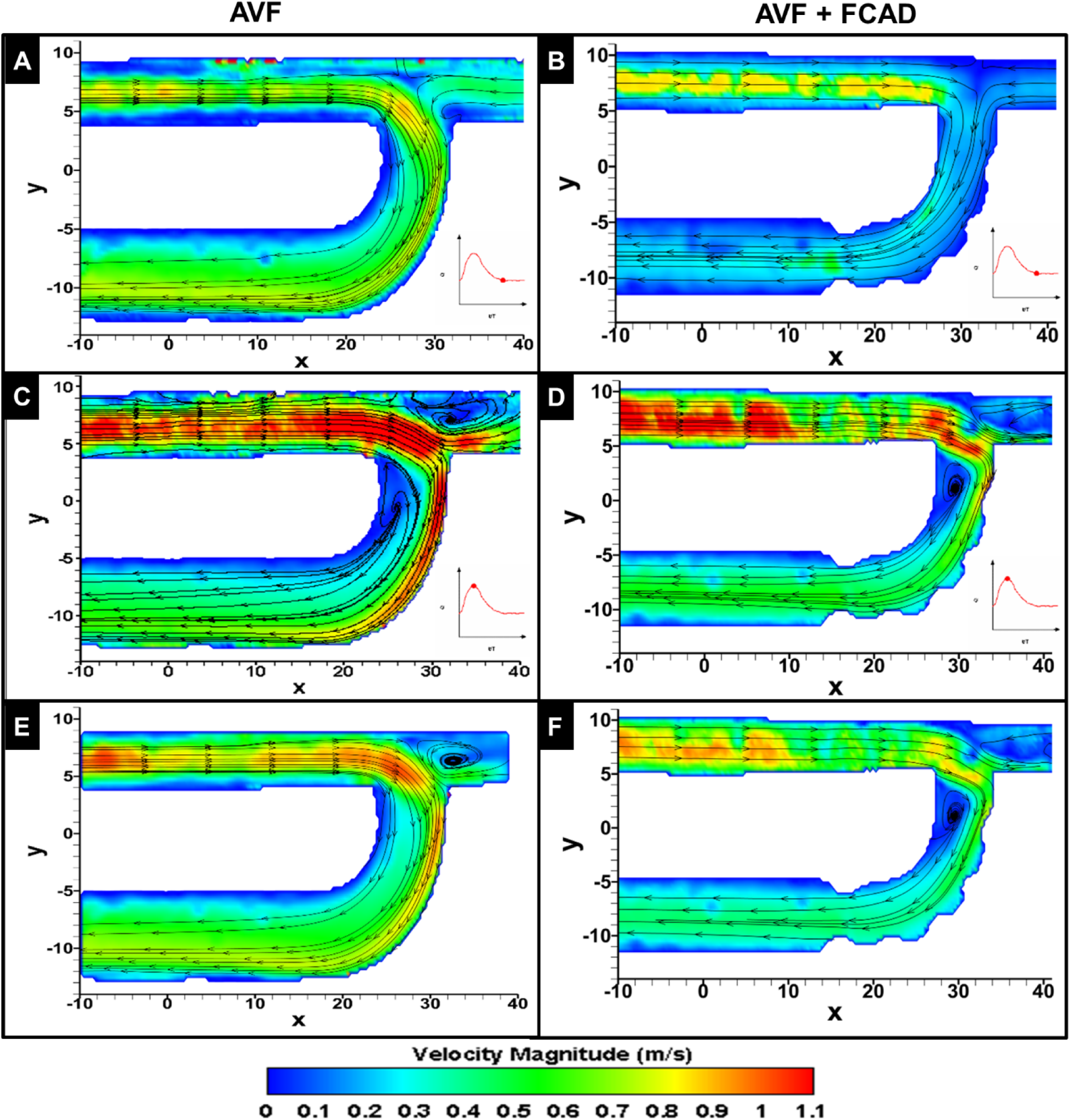
PIV-derived velocity flow fields in an idealized AVF and FCAD. Time-resolved PIV was used to quantitate the flow field in the anastomotic region of AVF phantoms with and without the FCAD. Temporal snapshots of velocity magnitude on the longitudinal symmetry plane are shown at mid-diastole (**A** & **B**), peak systole (**C** & **D**), and time-averaged across the entire flow cycle (**E** & **F**).

To further compare the flow field in the downstream vein, we quantified 2D axial velocity profiles along the mid-longitudinal plane at multiple diameters downstream of the venous outlet. As shown in Figure 8, flow in the AVF model was highly skewed toward the outer wall of the venous segment, even up to 3 vessel diameters (18 mm) downstream. In contrast, the axial velocity profile in the FCAD model was highly symmetric at the venous outlet, consistent with our CFD findings. Together, these findings validate the FCAD design’s ability to improve venous flow in the hemodynamic setting of an AVF.

**Figure 8:**
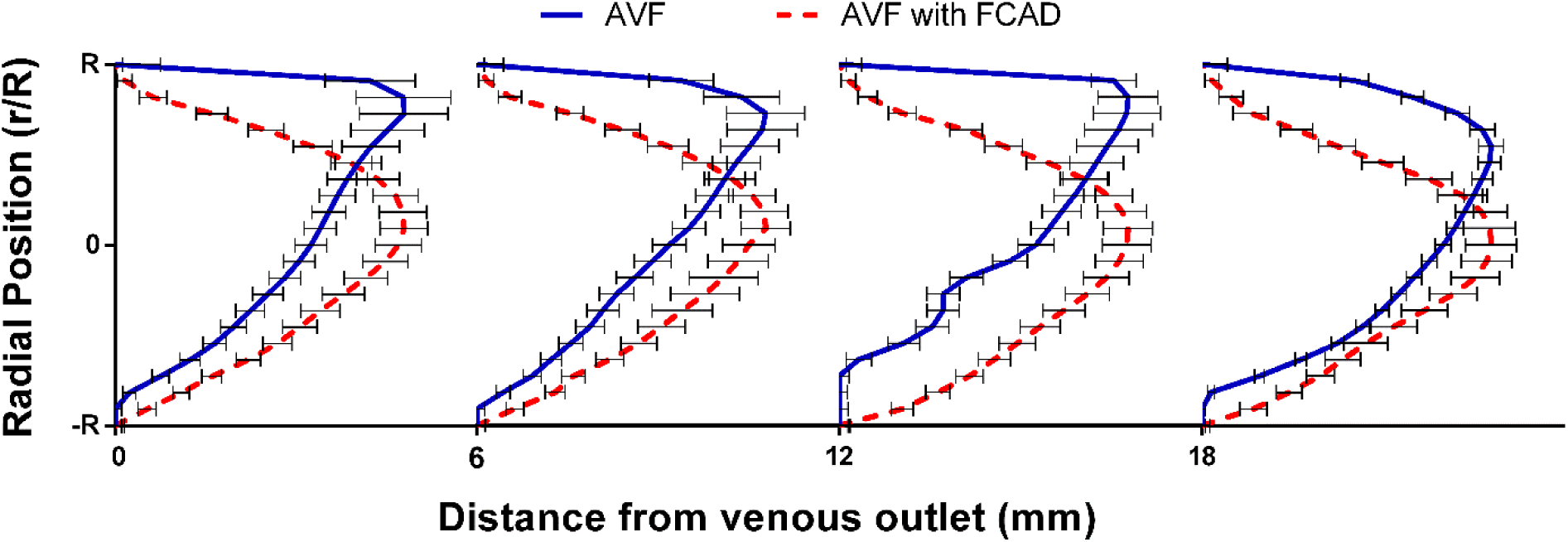
Comparison of PIV-derived axial velocity profiles in the AVF and FCAD models. Axial velocity measurements were recorded in the mid-longitudinal plane for each model and phase-averaged over 30 flow cycles. Each velocity measurement was normalized with respect to the radial position of the vessel (vessel axis = 0) and the maximum velocity. Error bars represent the root-mean-squared (RMS) error of each velocity measurement.

## Discussion

The recognition that disturbed flow affects endothelial function and vascular remodeling after AVF creation suggests that AVF maturation and patency may be improved by identifying an optimal AVF configuration. Several groups have investigated the influence of AVF configuration on WSS patterns and have suggested that small anastomotic angles may limit IH development.^16^ Furthermore, therapeutic approaches have sought to standardize AVF configuration, either endovascularly or extravascularly, to improve AVF flow and maturation. For example, the Optiflow device was introduced to maintain a fixed anastomotic angle, and its prosthetic material acts as a shield in the juxta-anastomotic region to prevent local stenosis.^17^ A clinical study of this device also confirmed its safety and surgical feasibility, with promising results.^18^ However, as concluded by Ene-Iordache et al., it is very difficult to identify an AVF configuration that completely abolishes all areas of disturbed WSS.^3^ In addition, the proximal venous segment and cephalic arch are the predominant sites of stenosis in brachiocephalic AVFs, suggesting that a greater vessel length is required for perturbations in flow to dissipate.^19^ This theory is further supported by findings from Mcgah et al. who demonstrated only a partial restoration of homeostatic WSS 20–30 mm away from the anastomosis in mature AVFs.^20^ Similarly, hemodynamic calculations have estimated that a vessel length exceeding 100 diameters is required to achieve settling of swirl and asymmetric flow.^21^ Therefore, we hypothesized that combining a “reasonable” configuration that shields the anastomotic region with a method to directly alter hemodynamic flow patterns in the draining vein could improve AVF outcomes.

Flow-conditioning technology is used in other applications to alter fluid flow patterns, yielding a reproducible downstream velocity profile over very short distances.^22^ By combining tab-style flow-conditioning technology with a method to standardize the AVF into a curved configuration, the FCAD has the potential to restore a physiological flow profile to the draining AVF vein. Incorporation of the FCAD geometry into a curved 90° AVF configuration demonstrated that the flow-conditioning tabs restore a symmetric, physiological flow profile at the venous outlet and minimize oscillatory WSS. In contrast, the reference AVF generated a large region of low flow along the inner venous wall, which was associated with an asymmetric flow profile and increased OSI in the downstream vein. Using PIV, we also demonstrate that similar flow-field phenomena occur in AVF flow phantoms with and without the FCAD. These results also align with those of Van Canneyt et al., who observed larger zones of low WSS in AVF with an anastomosis angle of 90°, but decreased vortex formation.^23^ A parametric analysis of the tab geometry also identified that tabs with an angle, height, and aspect ratio of 30°, 0.1 diameters, and 1.0, respectively, minimized regions of low and oscillatory WSS in the draining vein, two characteristics that have been shown to promote IH pathogenesis.^3^ However, increasing the tab height and aspect ratio were found to be inversely related to flow profile symmetry and to yield higher pressure losses. These results suggest the feasibility of the FCAD in effectively limiting negative vessel remodeling at the venous wall while shielding the juxta-anastomotic region from low oscillatory WSS.

To be effective, a flow conditioner should have low pressure losses, compact dimensions, low cost, and reduce flow disturbances, such as asymmetry and/or swirl. These design criteria align well with therapeutic approaches to vascular access, which are hindered by high costs and poor outcomes. In particular, pressure drop is a key metric, as it is an inverse measure of potential flow through an AVF and indicates whether adequate dialysis flow can be achieved. In our parametric analyses, we demonstrate that the incorporation of flow-conditioning tabs with small tab parameters not only improve venous flow but also impose minimal resistance to AVF flow across a series of flow regimes as measured by the pressure loss coefficient (ζ = 2.03, 2.04, 2.03, and 2.06 for AVF and tabs with angle, height, and aspect ratio of 30°, 0.1 diameters, and 1.0, respectively, at a Reynolds number of 1390). This FACD design corresponded to a mean pressure drop of 69 mmHg, which is consistent with the computed pressure drop previously reported by Hull et al. for a 90° end-to-side AVF with the same vessel sizes and mean flow rate.^24^ These computed pressure drops have also been shown to correlate well with in vivo measurements of pressure drop in brachiocephalic AVFs.^25^

Given the venous wall’s sensitivity to disturbed hemodynamics and injury, the introduction of prosthetic tabs that promote blunt-body turbulence and eddies into an autogenous AVF may be considered a drawback of this technique. However, trials investigating similar anastomotic devices, such as the Optiflow connector, have shown high degrees of safety following endovascular implantation of a device to standardize the AVF configuration^26^. Nonetheless, the impact of the flow alterations induced by the FCAD on turbulence generation and thrombogenesis warrants investigation. Using LNH to measure bulk fluid flow, we observed a significant change in flow direction upon contact with each tab; at the trailing edges, helical vortices with opposite directions to the bulk flow were generated. These counter-rotating vortices are thought to promote cross-stream mixing and induce a pressure gradient which pulls fluid towards the vessel wall to enhance the viscous boundary layer, resulting in the quick establishment of a homogeneous (i.e., conditioned) flow profile^5^. Further studies need to assess the fluid stresses induced by these vortices.

Although CFD has been used to evaluate pressure, flow, and WSS in idealized and patient-specific AVFs, the current study offers only a theoretical view of FCAD performance and has several limitations. First, the in silico models used to assess FCAD design and performance are based on an idealized AVF geometry and do not account for vessel wall compliance or changes in vessel size over time. While these assumptions are consistent with prior studies investigating WSS and flow metrics within an AVF, further fluid–structure interaction (FSI) analyses may be needed to assess the impact of FCAD and vessel elasticity on implant performance and durability. However, such FSI simulations do not significantly alter flow hemodynamics compared to rigid walls at the expense of a higher computational cost^27^. Second, although the peak-systolic Reynolds numbers calculated in the PA and venous outlet of the optimal FCAD geometry are relatively moderate (1265 and 2382, respectively), transitional and turbulent-like flow regimes with high-frequency fluctuations in stream-wise velocity have been observed in the juxta-anastomotic vein^13^. Therefore, additional studies are needed to determine whether transitional turbulent effects are present in our FCAD geometries and to validate the use of a laminar flow solver. Lastly, the presence of a transitional and turbulent flow regime may also lead to high shear loading on platelets traversing the endovascular device, leading to platelet activation and thrombosis^28^. Therefore, additional in-vitro and Lagrangian approaches for assessing platelet activation and thrombosis formation are being undertaken to minimize the risk of thrombosis in vivo^29^.

In summary, this study suggests the feasibility of the FCAD to normalize flow and WSS profiles in the draining AVF vein and identifies an ideal tab geometry with minimal resistance to AVF blood flow. Restoring physiological WSS levels on the venous wall is expected to preserve venous endothelial function and improve AVF maturation.

## Sources of Funding

This work was supported by a University of Cincinnati Technology Accelerator Award ODSA Tech 15-0160-1012977 (KS).

## Data Availability

The datasets generated and/or analyzed during the current study are not publicly available due to intellectual property considerations but are available from the corresponding author on reasonable request and with appropriate agreements in place.

## Acknowledgments

This work was supported in part by an allocation of computing time from the Ohio Supercomputer Center.

## Disclosures

Prabir Roy-Chaudhury is a Consultant/Advisor for WL Gore, Bard Peripheral Vascular, Medtronic, Cormedix, TVA, Akebia, Relypsa, and is the Founder/Chief Scientific Officer for Inovasc.

## Notes

### Competing Interest Statement

The authors have declared no competing interest.

## References

1. Cheung AK, Imrey PB, Alpers CE, et al. Intimal Hyperplasia, Stenosis, and Arteriovenous Fistula Maturation Failure in the Hemodialysis Fistula Maturation Study. J Am Soc Nephrol. 2017;28:1–9.

2. Badero OJ, Salifu MO, Wasse H, Work J. Frequency of swing-segment stenosis in referred dialysis patients with angiographically documented lesions. Am J Kidney Dis. 2008;51:93–98.

3. Ene-Iordache B, Remuzzi A. Disturbed flow in radial-cephalic arteriovenous fistulae for haemodialysis: low and oscillating shear stress locates the sites of stenosis. Nephrol Dial Transplant. 2012;27:358–368.

4. Franzoni M, Cattaneo I, Longaretti L, Figliuzzi M, Ene-Iordache B, Remuzzi A. Endothelial cell activation by hemodynamic shear stress derived from arteriovenous fistula for hemodialysis access. Am J Physiol - Hear Circ Physiol. 2016;310:H49–H59.

5. Laws EM, Harris R. Evaluation of a swirl-vor-tab flow conditioner. Flow Meas Instrum. 1993;4:101–108.

6. Frattolillo A, Massarotti N. Flow conditioners efficiency a comparison based on numerical approach. Flow Meas Instrum. 2002;13:1–11.

7. Remuzzi A, Ene-iordache B. Novel Paradigms for Dialysis Vascular Access : Upstream Hemodynamics and Vascular Remodeling in Dialysis Access Stenosis. 2013;8.

8. Krishnamoorthy MK, Banerjee RK, Wang Y, Choe AK, Rigger D, Roy-Chaudhury P. Anatomic configuration affects the flow rate and diameter of porcine arteriovenous fistulae. Kidney Int. 2012;81:745–750.

9. Saum K, Roy-Chaudhury P, Campos-Naciff B, Celdran-Bonafonte D. Arteriovenous Fistula Implant Effective for Inducing Laminar Blood Flow. *Int Pat Appl No US2016/049185, Publ No WO 2017/040366 A1*. 2017.

10. Division FE, Statement EP, Accuracy N, et al. Procedure for Estimation and Reporting of Uncertainty Due to Discretization in CFD Applications. J Fluids Eng. 2008;130:78001.

11. He Y, Terry CM, Nguyen C, Berceli SA, Shiu Y-TE, Cheung AK. Serial analysis of lumen geometry and hemodynamics in human arteriovenous fistula for hemodialysis using magnetic resonance imaging and computational fluid dynamics. J Biomech January. 2013;46:165–169.

12. Soulis J V., Lampri OP, Fytanidis DK, Giannoglou GD. Relative residence time and oscillatory shear index of non-Newtonian flow models in aorta. In: 10th International Workshop on Biomedical Engineering. ; 2011:1–4.

13. Bozzetto M, Ene-iordache B, Remuzzi A. Transitional Flow in the Venous Side of Patient-Specific Arteriovenous Fistulae for Hemodialysis Transitional Flow in the Venous Side of Patient-Specific Arteriovenous Fistulae for Hemodialysis. Ann Biomed Eng. 2016;44:2388–2401.

14. He X, Ku DN. Pulsatile flow in the human left coronary artery bifurcation: average conditions. J Biomech Eng. 1996;118:74–82.

15. Saggiomo V, Velders AH. Simple 3D Printed Scaffold-Removal Method for the Fabrication of Intricate Microfluidic Devices. Adv Sci. 2015;2:1–5.

16. Remuzzi A, Ene-Iordache B. Novel Paradigms for Dialysis Vascular Access: Upstream Hemodynamics and Vascular Remodeling in Dialysis Access Stenosis. Clin J Am Soc Nephrol. 2013;8:2186–2193.

17. Chemla E, Tavakoli A, Nikam M, et al. Arteriovenous fistula creation using the Optiflow^TM^ vascular anastomotic connector: The OPEN (Optiflow PatEncy and MaturatioN) study. J Vasc Access. 2014;15:38–44.

18. Chemla E, Velazquez CC, D’Abate F, Ramachandran V, Maytham G. Arteriovenous fistula construction with the VasQ^TM^ external support device: a pilot study. J Vasc Access. 2016;17:243–248.

19. Quencer K, Arici M. Arteriovenous Fistulas and Their Characteristic Sites of Stenosis. Vasc Interv Radiol. 2015;205:726–734.

20. McGah PM, Leotta DF, Beach KW, Eugene Zierler R, Aliseda A. Incomplete Restoration of Homeostatic Shear Stress Within Arteriovenous Fistulae. J Biomech Eng. 2012;135:11005.

21. Çarpınlıoğlu MÖ, Özahi E. Laminar flow control via utilization of pipe entrance inserts (a comment on entrance length concept). Flow Meas Instrum. 2011;22:165–174.

22. Ouazzane AK, Benhadj R. Flow conditioners design and their effects in reducing flow metering errors. Sens Rev Sens Rev Iss Multinatl Bus Rev. 2002;22:223–231.

23. Van Canneyt K, Pourchez T, Eloot S, et al. Hemodynamic impact of anastomosis size and angle in side-to-end arteriovenous fistulae: A computer analysis. J Vasc Access. 2010;11:52–58.

24. Hull J, Balakin B, Kellerman B, Wrolstad D. Computational fluid dynamic evaluation of the side-tp-side anastomosis for arteriovenous fistula. J Vasc Surg. 2013;58:187–193.

25. Browne L, Griffin P, Bashar K, Walsh S, Kavanagh E, Walsh M. In Vivo Validation of the In Silico Predicted Pressure Drop Across an Arteriovenous Fistula. Ann Biomed Eng. 2015;43:1275–1286.

26. Nikam M, Chemla ES, Evans J, et al. Prospective controlled pilot study of arteriovenous fistula placement using the novel Optiflow device. J Vasc Surg. 2015;61:1020–1025.

27. McGah PM, Leotta DF, Beach KW, Aliseda A. Effects of wall distensibility in hemodynamic simulations of an arteriovenous fistula. Biomech Model Mechanobiol. 2014;13:679–695.

28. Morbiducci U, Ponzini R, Nobili M, et al. Blood damage safety of prosthetic heart valves. Shear-induced platelet activation and local flow dynamics: a fluid-structure interaction approach. J Biomech. 2009;42:1952–1960.

29. Marom G, Bluestein D. Lagrangian methods for blood damage estimation in cardiovascular devices--How numerical implementation affects the results. Expert Rev Med Devices. 2016;13:113–122.

