## Supplemental Material for "Computational and Experimental Evaluation of a Flow-Conditioning Anastomotic Device for Arteriovenous Fistula Maturation"

#### Supplemental Materials

### Detailed Methods

##### *Idealized AVF Models with the FCAD*

Idealized models of an end-to-side brachiocephalic AVF with and without the FCAD implant were generated using SolidWorks 2015 (Dassault systems, Velizy-Villacoublay, France). We generalized the shape of the vessels, considering that every patient exhibits a unique vessel size and shape. The artery and vein was modeled as rigid circular cylinders with vessel diameters of 4.0 and 6.0 mm, respectively, and an anastomotic angle of 90°. The length of the proximal artery (PA), distal artery (DA), and vein were 12 times the vessel diameter in order to have sufficient hydraulic length to establish fully-developed flow and to characterize the flow field in the vein downstream of the anastomosis (Fig. 2A). The swing segment of the venous branch was generated with a radius of curvature equal to twice the venous diameter, and the juxta-anastomotic region of the vein was tapered for a length of two diameters to ensure a smooth transition between the artery and venous sections. For the FCAD implant models, we generated flow-conditioning tabs along the swing segment of the vein and outlet of the venous elbow (Fig. 2B). To characterize the impact of the tab geometry on the venous flow field, FCAD implant models were generated with four tab angles (15°, 30°, 45°, and 60°), four tab heights normalized to the venous diameter (0.1D, 0.2D, 0.3D, and 0.4D), and four tab aspect ratios (0.75, 1.0, 1.25, and 1.5) as shown in Figure 2C. A matrix of the FCAD models considered in the parametric analysis are shown in Table 1.

The fluid volume of each model was discretized with a structured hexahedral mesh generated in ICEM 16.0 (ANSYS, Inc., Canonsburg, PA). A boundary layer of 5 cells with a growth rate of 1.2 was generated at the walls to accurately capture changes in WSS. Three meshes of increasing element density were generated for a single model with and without the FCAD geometry to estimate the discretization error by the grid convergence index (GCI) method.[1] Steady-state computational fluid dynamics (CFD) simulations, corresponding to the venous flow rate at peak systole, were performed in each set of three meshes and used to estimate the mean velocity and mean WSS in the flow domain. The results of the grid convergence study are presented in Supplemental Table 1. Velocity and WSS fluctuations due to poor spatial resolution were minimized to less than 5% when the mesh size exceeded $5.4 \times 10^6$ and $7.0\times10^6$ elements in the AVF and FCAD model, respectively. Therefore, we choose to generate similar meshes (~$7.0 \times 10^6$) for all remaining FCAD models.

##### *CFD Simulations of Blood Flow*

Idealized AVF models were imported into the commercial CFD solver ANSYS Fluent 16.0 (Fluent, Inc., Lebanon, NH) and used to simulate pulsatile blood flow in the vessels. The transient Navier–Stokes equations were solved using a laminar model with a pressure-implicit with splitting of operators (PISO) algorithm for the pressure–velocity coupling. As boundary conditions, volumetric flow waveforms were prescribed at the PA and DA inlets (Fig. 2E), obtained from He et al. by 2D cine PC MRI of an end-to-side brachiocephalic AVF 5 months after creation.[2] A traction-free boundary condition was applied at the venous outlet, and zero velocity (no-slip condition) was applied at the vessel walls, which were considered to be rigid. Time-dependent terms were discretized in an implicit scheme with second-order accuracy. A total of 1,000 fixed time steps were used for each pulse cycle, resulting in a time step size of 0.001 seconds with a cardiac cycle period of 1 second. For each simulation, three complete cardiac cycles were solved to avoid start-up transients, and only the third cycle was saved for data processing in 100 equal time steps. A residual of $10^{-5}$ was set as the convergence criterion. Blood was modeled as an incompressible fluid with a constant density of 1.050 g/cm$^3$, and blood viscosity was considered non-Newtonian by using the Carreau rheological model, described previously as:

$$\mu = \mu_\infty + (\mu_0 - \mu_\infty)(1 + (\lambda\dot{\gamma})^2)^{\frac{n-1}{2}}$$

where $\mu_\infty$ is the limiting viscosity at infinite shear rate, $\mu_0$ is the limiting viscosity at zero shear rate, λ and n are constants, and $\dot{\gamma}$ is strain rate.[3] The use of the Carreau model was justified by our focus on the identification of venous wall regions experiencing low WSS, for which non-Newtonian effects may be particularly significant. For the PA inlet and venous outlet, we also calculated the Reynolds and the Womersley numbers as described previously.[4] Geometric and hemodynamic features of the AVF and FCAD model 2 simulations are summarized in Supplemental Table 2. The mean and range of the Reynolds numbers predicted at the vein outlet was 1315 (1137 – 2228) and 1643 (1343 - 2382) for the AVF and FCAD, respectively, justifying the use of a laminar flow solver.

To investigate the resistance to AVF flow imposed by implantation of the FCAD, we also performed CFD simulations in each AVF model by scaling the PA and DA waveforms to obtain a series of flow regimes with increasing Reynolds number. These studies were used to calculate the pressure loss coefficient of the FCAD across a range of flow regimes in which the device would have to function in vivo.

*Flow Field Characterization and Metric Analysis*

The ability of the FCAD to normalize venous AVF flow and WSS was investigated computationally by comparing flow profiles and WSS characteristics in the idealized AVF model (i.e., reference hemodynamic state) to the FCAD AVF models with parametric changes in their tab geometry. Once the flow field of each AVF was obtained, 13 transverse cross-sections were created normal to the centerline of the vein at 6 mm increments, starting at the anastomosis. The flow profile in each cross-section was assessed by means of velocity contours and the following absolute and relative efficiency parameters previously described in the literature.[5] By comparing these parameters at multiple locations downstream of the FCAD outlet to the corresponding location in the reference AVF model, we were able to tailor the performance of the device to improve venous flow through the AVF.

The profile symmetry, $K_{sym}$, was used to assess the symmetry of the time-averaged velocity profile with respect to the centerline of the vessel. This parameter represents a non-dimensional distance between the centroid of the flow profile and vessel axis and can be calculated in any transverse cross-section of the vessel and is expressed as:

$$K_{sym} = \frac{\sqrt{x_c^2 + y_c^2}}{R}$$

where R is the vessel radius, and $x_c$ and $y_c$ are the coordinates of the centroid of the mass flow given by:

$$x_c = \frac{\iint_A x\bar{v}dA}{Q} \qquad y_c = \frac{\iint_A y\bar{v}dA}{Q}$$

where v is the time-averaged axial velocity; Q the volumetric flow rate; and x and y are distances from the centerline to the radial coordinate. Therefore, the value of $K_{sym}$ is always a positive number, with smaller values indicating a less distorted flow.

Changes in WSS through the FCAD and downstream of the venous outlet were characterized using two WSS metrics to assess regions of potential endothelial dysfunction and/or IH formation. Time-averaged WSS (TAWSS) was used to identify regions of excessively high or low WSS which occurs over the cardiac cycle and was defined as:

$$TAWSS = \frac{1}{T}\int_0^T |\tau_w| dt$$

where T is the period of the cardiac cycle and $\tau_w$ is the instantaneous WSS vector. To quantify reciprocating disturbed flow during the cardiac cycle, the oscillatory shear index (OSI) was calculated on the surface of each AVF model as previously described[6]:

$$OSI = \frac{1}{2}\left(1 - \frac{\left|\int_0^T \tau_w dt\right|}{\int_0^T |\tau_w| dt}\right)$$

These WSS parameters were also averaged circumferentially in each of the 13 cross-sections, as previously described, resulting in a single value for each WSS parameter at multiple intervals along the AVF vessel.[2]

On the basis of the absolute parameters presented above, we also determined the relative efficiency of the FCAD at improving each flow-field metric, $\varepsilon_i$, compared to the reference AVF model as:

$$\varepsilon_i(z) = \frac{K_i^{AVF}(z) - K_i^{Device}(z)}{K_i^{AVF}(0) - K_i^{Device}(0)}$$

where $K^{AVF}$ and $K^{Device}$ represent the values of the absolute parameters calculated for the same vessel configuration without and with FCAD, respectively; and z is the distance downstream of the AVF anastomosis. Therefore, these parameters measure the efficiency of the FCAD compared to the reference AVF, as the distance downstream from the implant varies. Values of $\varepsilon_i$ greater than zero show an efficiency of the FCAD greater than the reference AVF and allow for comparison of different FCAD geometries.

To investigate vortex generation in the wake of the flow-conditioning tabs, the bulk flow phenotype was also characterized by determination of the local normalized helicity (LNH). As described by Levy et al, LNH is an effective tool for characterizing vortical structures past an axisymmetric body at a non-zero angle of attack and is expressed as[7]:

$$LNH = \frac{(\nabla \times v) \cdot v}{|\nabla \times v| \cdot |v|}$$

where $(\nabla \times v)$ and $v$ are the vorticity and the velocity vectors, respectively.

Lastly, we also assessed the resistance to AVF flow imposed by implantation of the FCAD. For such an implant to be clinically effective, its design must impose minimal resistance to venous flow through the AVF to not hinder blood flow to the hemodialysis machine. Therefore, we calculated the pressure loss coefficient (resistance coefficient), $\zeta$, for each FCAD model across a series of flow regimes with increasing Reynolds number by scaling the PA and DA flow waveforms (see section above). Flow rates were chosen to represent flows suitable for dialysis and potentially achievable with the vessel size modeled. The pressure loss coefficient is a dimensionless quantity representing pressure loss associated with the shape of pipes and obstacles to flow and was defined as[8]:

$$\zeta = \frac{P_{PA} - P_{venous\ outlet}}{0.5\rho V^2}$$

where V represents the mean flow velocity through the FCAD; ρ is the blood density; and $P_{PA}$ and $P_{venous\ outlet}$ are the total pressure at the PA and venous outlet, respectively. Pressures were measured 2.5mm from the anastomosis in the PA and the location corresponding to the venous outlet of the FCAD in all models. Compared with other measures of flow resistance, $\zeta$ characterizes both the energy losses due to viscous friction within the measurement region and the geometric shape of vessel. For these reasons, it was used to compare FCAD resistance between each model and the idealized AVF.

#### Figures and Tables

**Supplemental Table 1: Discretization error of mean velocity and wall shear stress**

| | AVF | | AVF with FCAD 2 | |
|---|---|---|---|---|
| | φ = mean velocity | φ = mean WSS | φ = mean velocity | φ = mean WSS |
| $N_1$ | 5,451,544 | 5,451,544 | 7,087,600 | 7,087,600 |
| $N_2$ | 2,467,070 | 2,467,070 | 3,169,331 | 3,169,331 |
| $N_3$ | 1,060,615 | 1,060,615 | 1,465,984 | 1,465,984 |
| $r_{21}$ | 1.303 | 1.303 | 0.110 | 0.110 |
| $r_{32}$ | 1.325 | 1.325 | 1.293 | 1.293 |
| $\varphi_1$ | 44.462 | 37.624 | 45.156 | 43.849 |
| $\varphi_2$ | 44.776 | 38.600 | 45.420 | 44.680 |
| $\varphi_3$ | 45.913 | 40.721 | 46.020 | 45.942 |
| p | 4.46 | 2.61 | 3.06 | 1.76 |
| $\varphi^{21}_{ext}$ | 44.322 | 36.641 | 44.949 | 42.470 |
| $e_a^{21}$ (%) | 0.7 | 2.6 | 0.6 | 1.9 |
| $e_{ext}^{21}$ (%) | 0.3 | 2.7 | 0.5 | 3.2 |
| $GCI^{21}_{fine}$ (%) | 0.4 | 3.3 | 0.6 | 3.9 |

The volume-averaged mean velocity was calculated for each at the maximum venous Reynolds number during peak systole (Re = 1775). The corresponding mean WSS was also calculated for each model. AVF = arteriovenous fistula, WSS = wall shear stress, GCI grid convergence index; for the other variables see Ref. 1.

**Supplemental Table 2: Geometric parameters and flow characteristics of CFD simulations**

| AVF model | AVF | AVF with FCAD 2 |
|---|---|---|
| N mesh cells | 5,451,544 | 7,087,600 |
| Cell vol ($x10^{-5}cm^3$) | 0.37(0.0064:1.19) | 0.29(0.0048:1.29) |
| N time steps/cycle | 1,000 | 1,000 |
| Delta t (s) | 0.001 | 0.001 |
| Q PA (mL/min) | 705(563:984) | 705(563:984) |
| Q DA (mL/min) | -22(-230:62) | -22(-230:62) |
| Q PA:DA:V (%) | 100:23:77 – 91:9:100 | 100:23:77 – 91:9:100 |
| Re PA | 906(723:1265) | 906(723:1265) |
| α PA | 3.43 | 3.43 |
| Re Venous Outlet | 1315(1137:2228) | 1643(1343:2382) |
| Mean Pressure Drop (mm Hg) | 66(52:78) | 69(53:81) |

Delta t, blood flow rates, Reynolds number (Re), and pressure drop are expressed as time-averaged and (minimum:maximum) values over the pulse cycle. Q PA:DA:V represents the blood flow division ratio between these limbs of the AVF throughout the cardiac cycle. Cell vol represents the volume of the cells of the mesh expressed as mean(minimum:maximum). Q, volumetric blood flow rate; PA, proximal artery; DA, distal artery; V, vein; Re, Reynolds number; α, Womersley number.

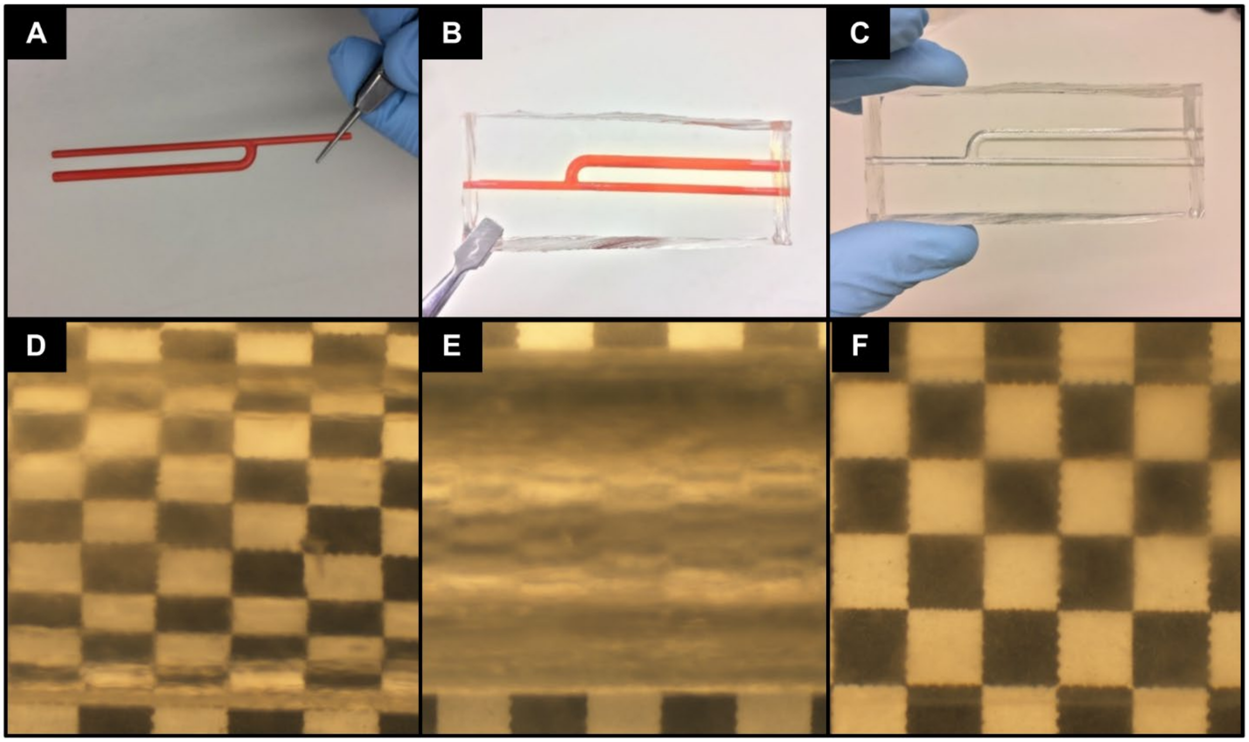


**Supplemental Figure 1**: Fabrication of AVF flow phantoms for PIV studies. [A] Example of 3D printed AVF model at at a 4:1 scale. [B-C] Pictures of the fabrication process after imbedding the model in PDMS (B) and subsequently dissolving the model to produce a hollow phantom (C). [D-E] Refractive index matching tests for an AVF phantom filled with (D) 100 % water, (E) air, and (F) a refractive index-matched working fluid of glycerol and water. Note the absence of refraction in the background tile pattern with the index-matched fluid compared to water and air.

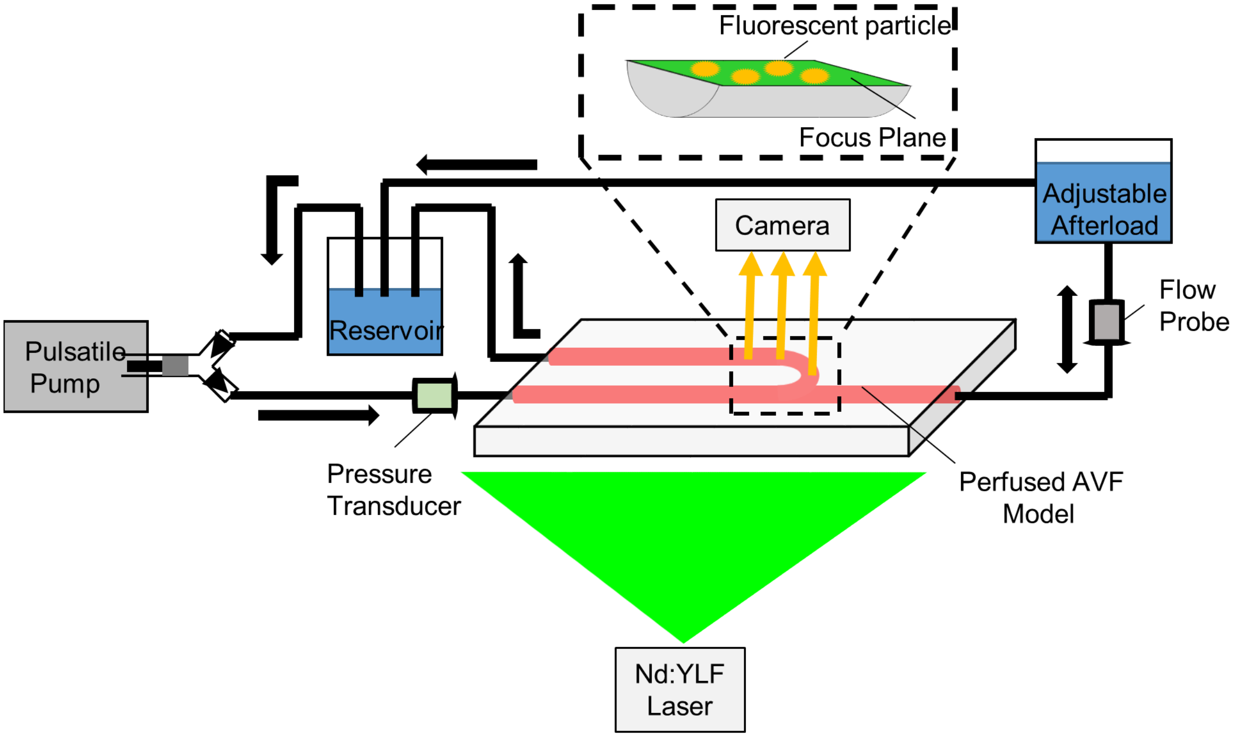


**Supplemental Figure 2:** Schematic of in vitro flow system for PIV studies. A working fluid containing fluorescent particles was perfused with a pulsatile blood pump through the AVF flow phantoms. PIV imaging was performed at the anastomosis and venous outlet using a Nd:YFL laser to excite the particles, and a high-speed camera to record particle motion.

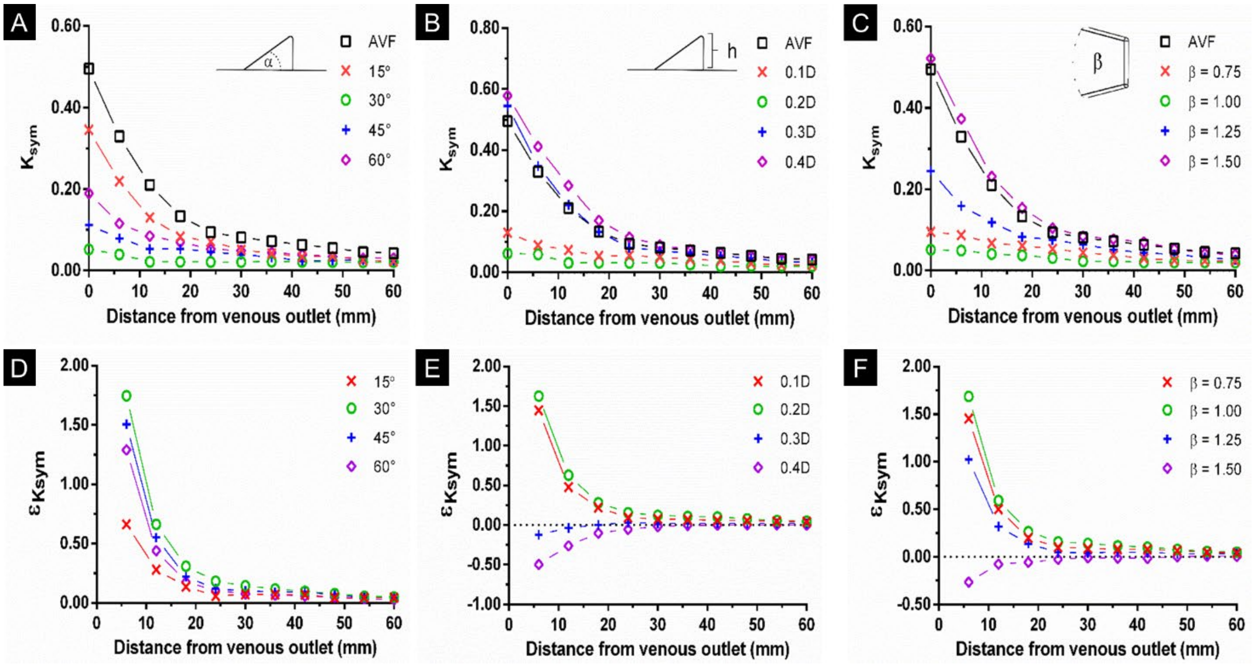


**Supplemental Figure 3:** Parametric analysis of flow field symmetry along the draining vein in AVF and FCAD models. Symmetry number, Ksym, was calculated at multiple cross-sections downstream of the anastomosis in FCADs with increasing [**A**] tab angle (α), [**B**] tab height, and [**C**] tab surface area ratio. The location of the FCAD venous outlet was set to be 0, and the anastomosis is labeled "A" in each plot axis. The reference AVF without an FCAD is labeled "AVF". The relative efficiency (ε) of each FCAD at improving the symmetry number, εKsym, compared to the reference AVF was also determined for each tab angle (**D**), height (**E**), and area ratio (**F**).

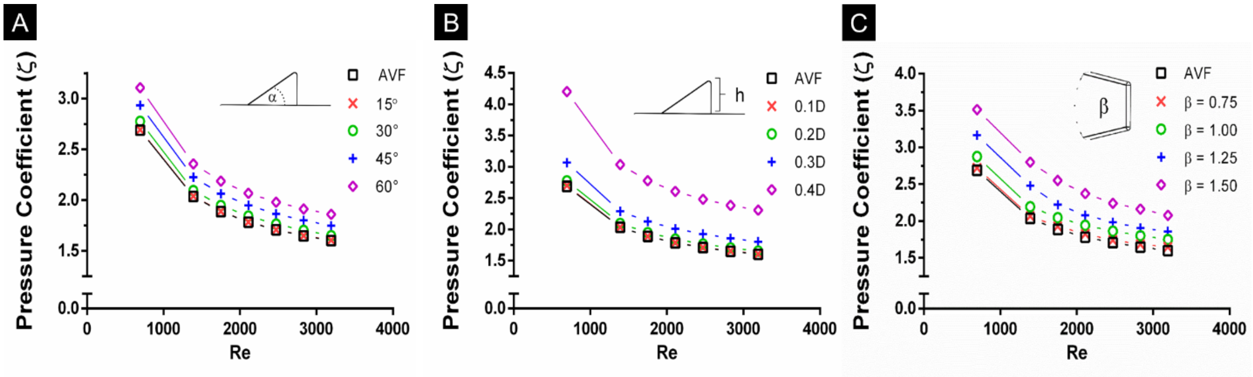


**Supplemental Figure 4:** Parametric analysis of pressure drop in AVF and FCAD models. The pressure loss coefficient, ζ, was calculated for FCADs with increasing [**A**] tab angle, [**B**] tab height, and [**C**] tab surface area ratio across a series of flow regimes with increasing Reynolds number (Re). The reference AVF without an FCAD is also shown.

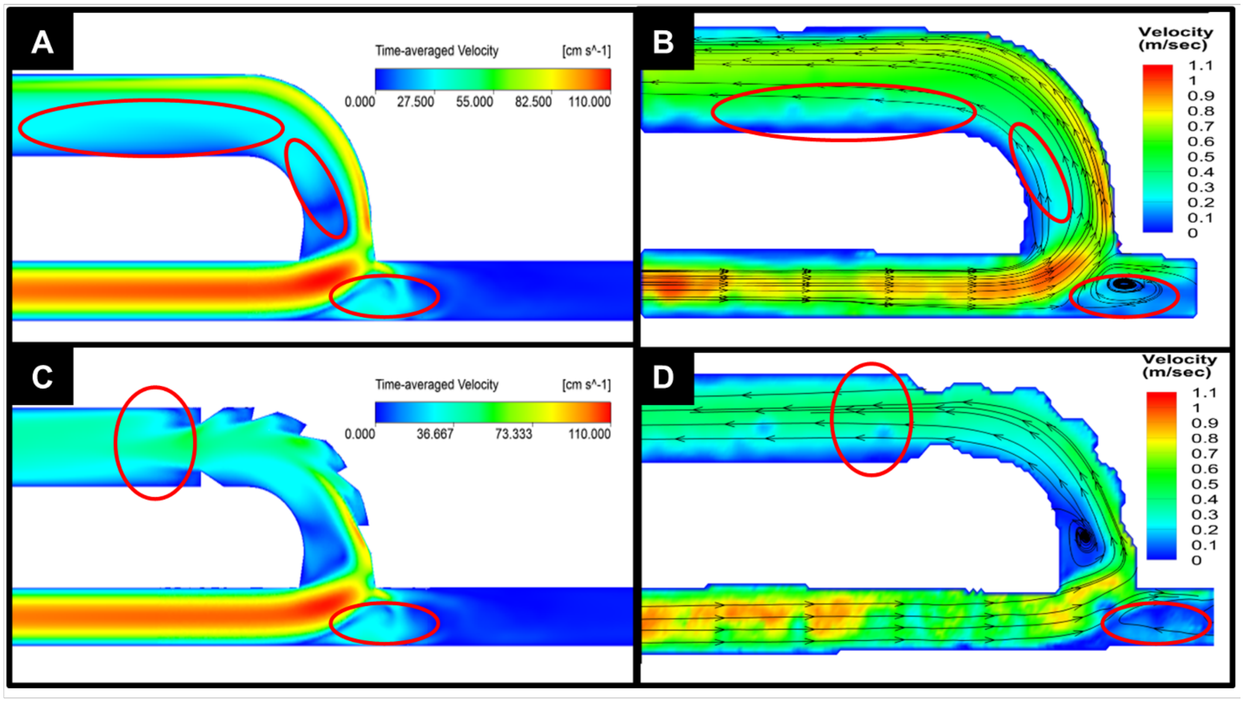


**Supplemental Figure 5:** Velocity field comparison between the PIV experimental results and the CFD simulations. The circled red regions show similar time-averaged flow features between the CFD and PIV results in both the AVF model [**A & B**] and FCAD [**C & D**] models. All pictures are on the same color scale shown in the CFD images.

#### References


1. Division FE, Statement EP, Accuracy N, et al. Procedure for Estimation and Reporting of Uncertainty Due to Discretization in CFD Applications. *J Fluids Eng*. 2008;130:78001.
2. He Y, Terry CM, Nguyen C, Berceli SA, Shiu Y-TE, Cheung AK. Serial analysis of lumen geometry and hemodynamics in human arteriovenous fistula for hemodialysis using magnetic resonance imaging and computational fluid dynamics. *J Biomech January*. 2013;4.
3. Soulis J V., Lampri OP, Fytanidis DK, Giannoglou GD. Relative residence time and oscillatory shear index of non-Newtonian flow models in aorta. In: *10th International Workshop on Biomedical Engineering, BioEng 2011*. ; 2011.
4. Bozzetto M, Ene-iordache B, Remuzzi A. Transitional Flow in the Venous Side of Patient-Specific Arteriovenous Fistulae for Hemodialysis Transitional Flow in the Venous Side of Patient-Specific Arteriovenous Fistulae for Hemodialysis. *Ann Biomed Eng*. 2016;44:2388-2401.
5. Frattolillo A, Massarotti N. Flow conditioners efficiency a comparison based on numerical approach. *Flow Meas Instrum*. 2002;13:1-11.
6. He X, Ku DN. Pulsatile flow in the human left coronary artery bifurcation: average conditions. *J Biomech Eng*. 1996;118:74-82.
7. Levy Y, Deganif D, Seginer A. Graphical Visualization of Vortical Flows by Means of Helicity. *AIAA J*. 1990;28:1347-1352.
8. Çarpınlıoğlu MÖ, Özahi E. Laminar flow control via utilization of pipe entrance inserts (a comment on entrance length concept). *Flow Meas Instrum*. 2011;22:165-174.
